# A molecularly defined brain circuit module for regulating the panic-like defensive state

**DOI:** 10.1101/2024.12.20.629854

**Authors:** Miao Zhao, Li Zhang, Zhenhua Chen, Shuangfeng Zhang, Xinyu Cheng, Meizhu Huang, Xiating Li, Huating Gu, Xuyan Guan, Dandan Geng, Yaning Li, Yiheng Tu, Zhiyong Xie, Fan Zhang, Huijie Ma, Dapeng Li, Qingfeng Wu, Peng Cao

**Author notes:** These authors contributed equally (Miao Zhao, Li Zhang, Zhenhua Chen, Shuangfeng Zhang, Xinyu Cheng). Correspondence (P.C.), (Q.W.), (D.L.).

## Abstract

Panic is an episode of strong defensive state, characterized by intense fear and severe physical symptoms such as elevated cardiorespiratory activities. How the brain generates panic state remains poorly understood. Here, we developed a robot-based experimental paradigm to evoke panic-like defensive state in mice. When stimulated by the robot, the mice exhibited jumping escapes and elevated cardiorespiratory activities. With this paradigm, we identified *Cbln2*-expressing (*Cbln2*+) neurons in the posterior hypothalamic nucleus (PHN) as a key neuronal population essential for the induction of panic-like defensive state. Activation of *Cbln2*+ PHN neurons induced behavioral and physical symptoms of panic-like defensive state. These neurons were strongly activated by noxious mechanical stimuli and encode jumping escape vigor. They were synaptically innervated by anxiety-associated brain areas and provoked panic-like defensive state via their projection to the periaqueductal gray. Together, our results reveal a molecularly defined circuit module that regulates the panic-like defensive state in mice.

## INTRODUCTION

Panic is an episode of strong defensive state with important biological functions and psychiatric relevance. In biology, panic state is provoked by imminent life-threatening danger (<u>Stein and Bouwer, 1997</u>; <u>McNaughton and Corr, 2004</u>; <u>Hoffman et al., 2022</u>). During panic state, the body initiates both physiological (e.g., elevated cardiorespiratory activities) and behavioral responses (e.g., fight or flight), thus facilitating escape from dangerous situations (<u>Griebel et al., 1996</u>; <u>Blanchard et al., 2001</u>). In psychiatry, panic disorder is a type of anxiety disorder characterized by unexpected and recurrent episodes of panic state, known as panic attacks (<u>American Psychiatric Association, 2013</u>). During panic attack, patients exhibit a series of symptoms, including tachycardia, tachypnea, and an intense desire to escape (<u>American Psychiatric Association, 2013</u>), mirroring the defensive responses observed in biological panic state. Panic disorder affects up to 5% of the population at some stage in their lives (<u>Kessler et al., 2006</u>). Given the biological importance and psychiatric relevance, it is critical to understand how the brain generates the panic state.

In recent decades, the neurobiology of panic state has been intensively studied in human subjects and animal models (for reviews, see <u>Graeff and Del-Ben, 2008</u>; <u>Johnson et al., 2014</u>; <u>Guan and Cao, 2023</u>), yielding three lines of important findings. <u>First</u>, considerable efforts have concentrated on delineating the neuroanatomical basis of panic disorder (<u>Gorman et al., 2000</u>; <u>Dresler et al., 2013</u>). Notably, deep brain stimulation in the posterior hypothalamus (<u>Schoenen et al., 2005</u>; <u>Bartsch et al., 2008</u>) and periaqueductal gray (PAG) (<u>Nashold et al., 1969</u>; <u>Del-Ben and Graeff, 2009</u>) induced panic attacks in human. Analogous pharmacological stimulations of these brain regions in rats also elicited panic-like defensive responses (<u>Shekhar and DiMicco, 1987</u>; <u>Jenck et al., 1995</u>), suggesting an important role of these brain areas in the generation of panic state. <u>Second</u>, pharmacological studies have revealed a series of neurotransmitters or neuromodulators implicated in panic disorder, including serotonin (5-HT) (<u>Graeff et al., 1996</u>; <u>Maron and Shlik, 2006</u>), GABA (<u>Zwanzger and Rupprecht, 2005</u>), and orexin (<u>Johnson et al., 2012</u>). Selective serotonin reuptake inhibitors (e.g., fluoxetine) and benzodiazepines are effective in treating panic disorder (<u>Quagliato et al., 2018</u>). <u>Third</u>, panic attacks can be evoked by mild interoceptive stimuli in patients with panic disorder (<u>Domschke et al., 2010</u>; <u>Hamm et al., 2014</u>). Inhalation of carbon dioxide (CO_2_) can induce panic attack in panic disorder patients (<u>Gorman et al., 1994</u>) and panic-like responses in animal models (<u>Leibold et al., 2016</u>), suggesting chemosensation of acidosis may trigger false suffocation alarms associated with panic attack (<u>Klein, 1993</u>; <u>Wemmie, 2011</u>). In addition, lactate administration can provoke panic attack (<u>Liebowitz et al., 1984</u>), which may depend on neural pathways mediated by circumventricular organs (<u>Shekhar and Keim, 1997</u>).

Despite the above discoveries, the synaptic and circuit mechanisms in the brain that underlie the generation of panic state remain elusive, leaving two important questions unanswered. <u>First</u>, the key neuronal populations responsible for provoking panic state has not been identified. <u>Second</u>, how the panic-provoking neurons are synaptically connected to other panic-associated brain areas is uncharted.

To address the aforementioned questions, it is essential to establish an appropriate experimental paradigm for panic induction. Rats have been extensively used for modeling panic disorder (<u>Beckett et al., 1996</u>; <u>Graeff et al., 1998</u>; <u>Johnson and Shekhar 2012</u>; <u>Moreira et al., 2013</u>). For example, chronic inhibition of GABA synthesis in the dorsomedial/perifornical hypothalamic region of rats induces panic vulnerability to mild interoceptive stimuli, providing an excellent model with robust face, predictive, and construct validity (<u>Johnson and Shekhar, 2012</u>). Nevertheless, the limited capacity of rats for genetic manipulation, which is crucial for studying molecularly defined synaptic and circuit mechanisms underlying panicogenesis, presents a challenge. Griebel et al. (1996) used predator-induced flight responses in mice as an experimental model of panic attacks. However, this paradigm requires an experimenter to hold an anesthetized rat as a predator, thus introducing human interventions into mouse behavior. Optimizing this paradigm is a critical step towards exploring the synaptic and circuit mechanisms underlying panic-like defensive state.

In this study, we developed a robot-based paradigm to induce panic-like defensive state in mice. In this paradigm, mice exhibited explosive jumping escapes, tachycardia, and tachypnea, all of which can be quantitatively measured. Using this paradigm, we identified *Cbln2*-expressing (*Cbln2*+) neurons in the posterior hypothalamic nucleus (PHN) as a key neuronal population essential for panic-like state. Optogenetic activation of *Cbln2*+ PHN neurons evoked jumping escapes, tachycardia, and tachypnea, thus mimicking panic-like defensive state. These neurons responded to noxious mechanical stimuli and encoded flight response vigor. They were innervated by multiple panic-associated brain areas and provoked panic-like defensive state via projection to the PAG. Together, these results reveal a molecularly defined circuit module that regulates panic-like defensive state in mice.

## RESULTS

### Experimental paradigm to elicit panic-like defensive state in mice

In this study, we developed an experimental paradigm to evoke a panic-like defensive state in mice. As a mouse navigated an arena (15 × 15 cm), we introduced an electrically powered beetle-like robot (4 × 6 × 8 cm, 23 g) that moved at a constant speed (30 cm/s). Collisions with the robot prompted the mouse to exhibit explosive jumping escapes (<u>Supplementary Video 1</u>). To quantitatively characterize the defensive state of mice, we recorded their jumping escape, electrocardiogram (ECG), and breathing (Fig. 1, a & b) (<u>Grimaud and Murthy, 2018</u>; <u>Signoret-Genest et al., 2023</u>). Quantitative analyses indicated that, following the introduction of the robot, the mice exhibited frequent jumping escapes (<u>Supplementary Video 2</u>; Supplementary Fig. 1, a-c) in parallel with persistent increases in both heart (<u>Supplementary Video 3</u>; Supplementary Fig. 1, d-f) and breathing rates (<u>Supplementary Video 4</u>; Supplementary Fig. 1, g-i). Fluoxetine has been used to alleviate panic state in patients (<u>Quagliato et al., 2018</u>). We found that chronic treatment with the panicolytic drug fluoxetine (21 days, 5 mg/kg per day) significantly reduced these robot-induced defensive responses (Fig. 1<u>, c & d</u>), suggesting that the observed behavioral and autonomic responses are indicative of a panic-like defensive state.

**Fig. 1.**
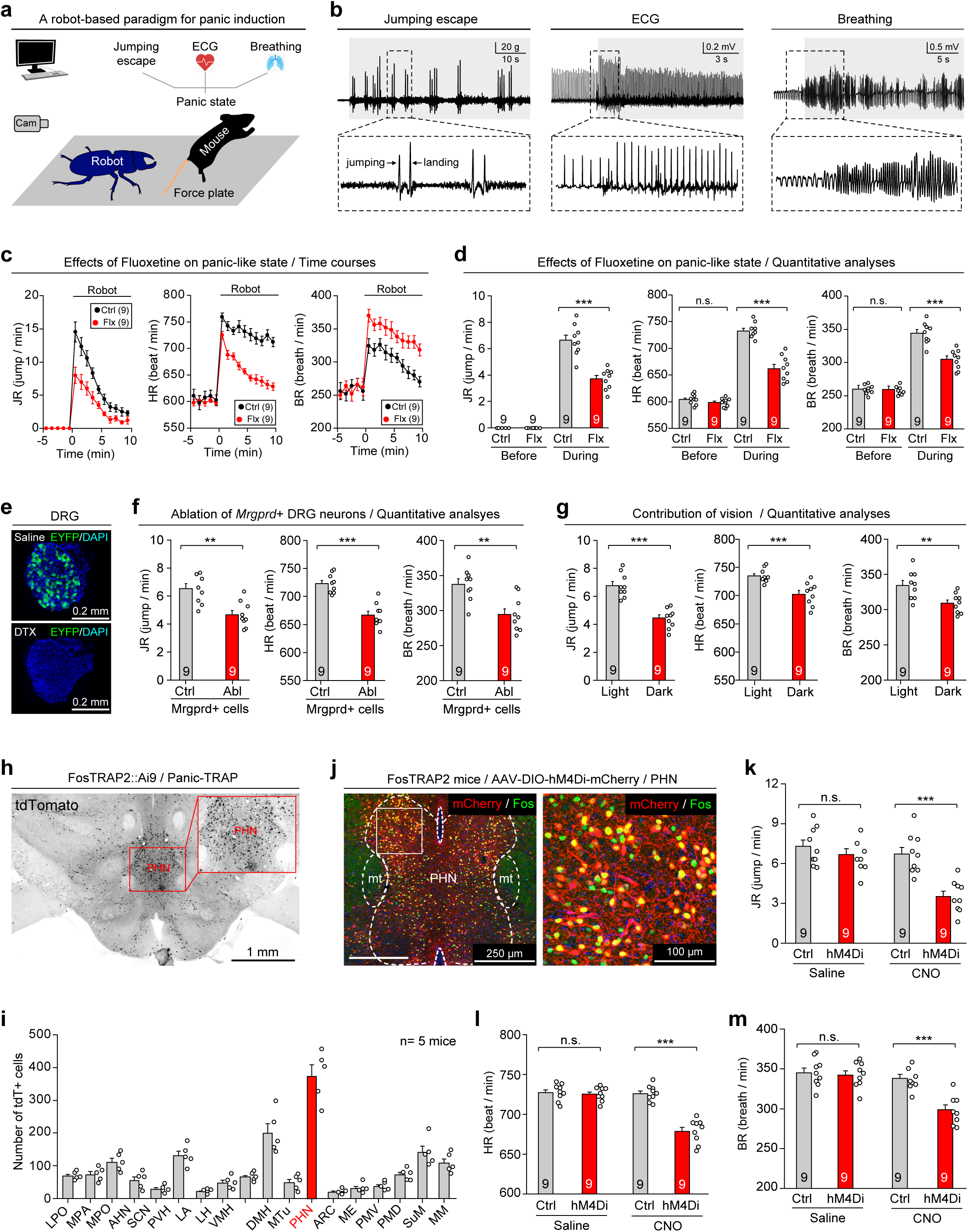
Identification of the PHN as a key brain area for panic-like defensive state. **(a)** Schematic diagram showing experimental paradigm for measuring robot-induced panic-like defensive responses. (**b**) Example traces showing jumping escape (*left*), ECG (*middle*), and breathing (*right*) before and during panic induction. Jumping escape was identified as a pair of force-plate pulses indicating jumping and landing, respectively. **(c, d)** Time courses (c) and quantitative analyses (d) of jumping rate (JR), heart rate (HR), and breathing rate (BR) before and during panic induction in mice without (Ctrl) or with chronic fluoxetine treatment (Flx). **(e)** Example micrographs showing ablation of *Mrgprd*+ neurons (green) in DRG. **(f)** Quantitative analyses of jumping rate (JR), heart rate (HR), and breathing rate (BR) during panic induction in mice with and without ablation of *Mrgprd*+ DRG neurons. **(g)** Quantitative analyses of jumping rate (JR), heart rate (HR), and breathing rate (BR) during panic induction in mice with and without light in the arena. **(h)** Example coronal brain section of FosTRAP2::Ai9 mouse showing panic-associated neurons labeled with tdTomato (gray-scale). **(i)** Quantitative analyses of tdTomato-labeled neurons in different hypothalamic areas of mice subjected to panic induction. **(j)** Example coronal brain section (*left*) and magnified fields (*right*) showing hM4Di-mCherry expression induced by Panic-TRAP experiment and Fos expression induced by Panic-Fos experiment. **(k-m)** Quantitative analyses of jumping rate (k), heart rate (l), and breathing rate (m) during panic induction in mice treated with saline or CNO (1 mg/kg) to chemogenetically inactivate Panic-TRAPed PHN neurons. Scale bars are labeled in graphs. Numbers of mice (c, d, f, g, i, k-m) are indicated in graphs. Data in (c, d, f, g, i, k-m) are means ± standard error of the mean (SEM). Statistical analyses (d, f, g, k-m) were performed using Student *t-*test (*** *P* < 0.001; \*\**P* < 0.01; \**P* < 0.05; n.s. *P* > 0.1). For *P*-values, see Supplementary Table 3.

We then investigated the involvement of sensory systems in the robot-induced panic-like defensive state. *Mrgprd*+ sensory neurons in the dorsal root ganglion (DRG) are a subset of non-peptidergic sensory neurons that respond to noxious mechanical stimuli (<u>Cavanaugh et al., 2009</u>). In Mrgprd-CreERT2/iDTR/Ai3 mice (<u>Olson et al., 2017</u>; <u>Buch et al., 2005</u>; <u>Madisen et al., 2010</u>), tamoxifen-induced DTR expression and subsequent treatment of diphtheria toxin resulted in a decrease (88% ± 6%, n = 9 pairs) in the number of EYFP+ DRG neurons (putatively *Mrgprd*+) (Fig. 1e). Ablation of *Mrgprd*+ DRG neurons significantly impaired robot-induced defensive responses in mice (Fig. 1f; Supplementary Fig. 1j), suggesting that noxious mechanical stimuli during robot-mouse collisions may play an important role in panic induction in this paradigm. Conducting the experiment in darkness (∼0.002 lux) significantly reduced the robot-induced defensive responses (Fig. 1g; Supplementary Fig. 1k), implying a potential role of visual detection of impending collision in eliciting a panic-like defensive state in this paradigm. These results suggest that the robot-mouse collision is a key factor to induce the panic-like defensive state. Thus, we employed this robot-based paradigm to explore the synaptic and circuit mechanisms underpinning the panic-like defensive state in mice.

### Importance of PHN neurons in panic-like defensive state

We next performed whole-brain mapping of neurons associated with panic-like defensive state by using the FosTRAP2 strategy. FosTRAP2 (Fos-2A-CreERT2) was designed to express a tamoxifen-inducible Cre recombinase in neurons that are activated during specific behaviors (<u>Allen et al., 2017</u>). To genetically label panic-associated neurons in mouse brain, we intraperitoneally injected 4-hydroxy-tamoxifen (4-OHT, 30 mg/kg) into FosTRAP2::Ai9 mice and then induced the panic-like defensive state using the robot-based paradigm (Supplementary Fig. 2a). This Panic-TRAP approach resulted in the labeling of abundant tdTomato+ neurons in the brain (Supplementary Fig. 2b); in contrast, Ctrl-TRAP mice, which received 4-OHT without exposure to the robot, showed few tdTomato-labeled neurons in the brain (Supplementary Fig. 2c). Quantification of Panic-TRAPed neurons (tdTomato+) in various hypothalamic nuclei (<u>Paxinos and Franklin, 2008</u>) revealed the highest number in the PHN (Fig. 1<u>, h & i</u>). Based on this morphological observation, we hypothesized that PHN neurons may be important for the panic-like defensive state in mice.

To explore the functional significance of PHN neurons in the panic-like defensive state, AAV-DIO-hM4Di-mCherry was injected into the PHN of FosTRAP2 mice (Supplementary Fig. 2d). In Ctrl-TRAP mice without panic induction, few PHN neurons were labeled (228 ± 24, n = 9 mice); however, in Panic-TRAP mice, abundant PHN neurons were labeled with hM4Di-mCherry (6,620 ± 326 cells, n = 9 mice) (Supplementary Fig. 2, e & f). To examine the specificity and efficiency of hM4Di-mCherry labeling, Fos and mCherry in the PHN of Panic-TRAP mice were immunostained (Fig. 1j; Supplementary Fig. 2g). Quantitative analyses indicated that a large proportion of mCherry+ cells (82% ± 6%, n = 5 mice) were positive for Fos, and most Fos+ cells (79% ± 4%, n = 5 mice) were positive for mCherry (Supplementary Fig. 2h). Chemogenetic inactivation of these neurons with intraperitoneal administration of clozapine N-oxide (CNO, 1 mg/kg) resulted in a notable reduction in robot-induced jumping escapes (Fig. 1k; Supplementary Fig. 2i), heart-rate increase (Fig. 1l; Supplementary Fig. 2j) and breathing-rate increase (Fig. 1m; Supplementary Fig. 2k). These findings suggest the PHN neurons are essential for inducing the panic-like defensive state in mice.

### Molecular identities of panic-associated PHN neurons

Next, we examined the molecular identities of PHN neurons that are involved in the panic-like defensive state in mice. Two weeks after Panic-TRAP experiment, the PHN and the adjacent brain areas of FosTRAP2::Ai9 mice were micro-dissected for single-nucleus RNA-seq (snRNA-seq) (Fig. 2a). A total of 42,926 qualified cells from three batches of brain tissues were captured. They were categorized into seven major cell classes that expressed well-established marker genes for neurons, microglia, ependymal cells, oligodendrocytes, astrocytes, oligodendrocyte precursor cells, and endothelial cells (Fig. 2, b & c; Supplementary Fig. 3, a & b; Supplementary Table 4). Unsupervised graph-based clustering of 29,255 neurons revealed thirteen neuronal clusters that express specific molecular markers (Fig. 2<u>, d & e</u>; Supplementary Table 5). By cross-referencing the Allen Mouse Brain Atlas (https://mouse.brain-map.org), we found that six neuronal clusters (N1-N6) were distributed within the PHN (Supplementary Fig. 3, c-h), whereas the other seven clusters (N7-N13) were in brain areas that are adjacent to the PHN (Supplementary Fig. 3, i-o). We calculated the expression level of different marker genes and identified the top markers that optimally label the six neuronal clusters in the PHN, including *Cbln2* for N1, *Slc32a1* for N2, *Crhbp* for N3, *Slc17a8* for N4, *Tac1* for N5, and *Sst* for N6 (Fig. 2<u>, f-k</u>). We also examined the expression of molecular markers for glutamatergic (*Slc17a6*) and GABAergic (*Slc32a1*) neurons in these six clusters, finding that N2 were GABAergic and the other five clusters were glutamatergic (Fig. 2<u>, l & m</u>).

**Fig. 2.**
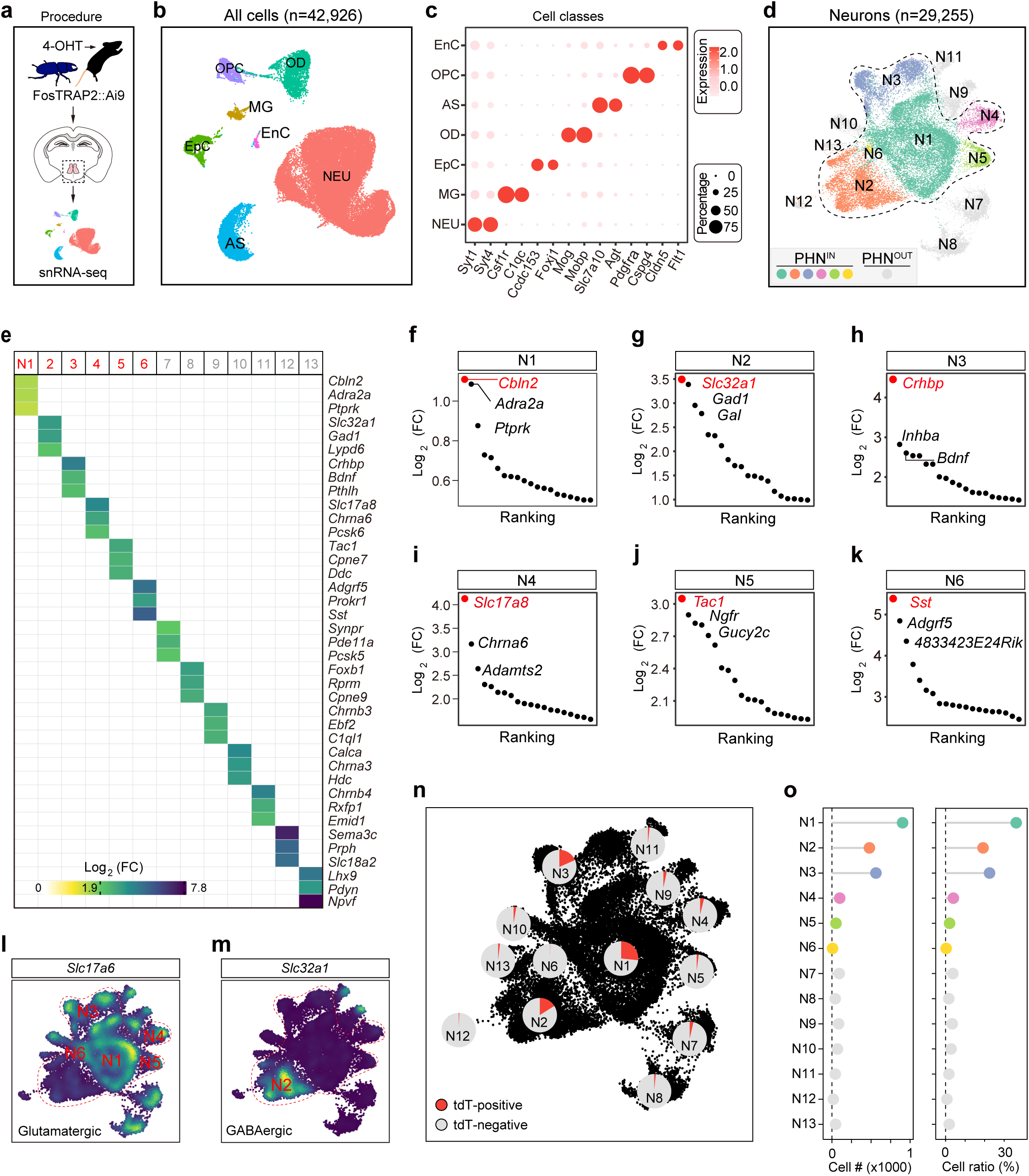
Molecular profiling of PHN neurons and genetic identification of panic-TRAP neuronal subtypes. **(a)** Schematic diagram showing the procedure of Panic-TRAP experiment to label panic-associated PHN neurons and subsequent transcriptomic profiling of these PHN neurons at single-nucleus resolution. Adult FosTRAP2::Ai9 mice were pre-treated with 4-OHT and subjected to robot-based panic induction. Two weeks after panic-TRAP, brain tissues containing the PHN were micro-dissected for snRNA-seq. **(b)** UMAP visualization of single-nucleus transcriptomic dataset containing 42,926 hypothalamic PHN cells collected from Fos-TRAP2::Ai9 mice (n= 15 mice). Cell classes included astrocytes (AS), endothelial cells (EnC), microglia (MG), neurons (NEU), oligodendrocyte precursor cells (OPC), oligodendrocytes (OD), and ependymal cells (EpC). **(c)** Dot plot showing relative expression levels of markers in different cell types. **(d)** UMAP visualization and sub-clustering of 29,255 neurons, with neuronal clusters beyond PHN (N7-N13) shown in gray. **(e)** Heat-map showing normalized expression of marker genes for each neuronal cluster. neuronal clusters in the PHN (N1-N6) are highlighted in red. FC, fold-change. **(f-k)** Dot plots showing ordering of marker genes in PHN neuronal clusters (N1-N6), with top markers in red. **(l, m)** Feature plots illustrating expression patterns of *Slc17a6* (l) and *Slc32a1* (m) in different PHN neuronal clusters. **(n)** Pie chart showing percentage of Panic-TRAPed neurons (tdTomato+) in each neuronal cluster. **(o)** Lollipop charts presenting cell numbers (*left*) and cell ratio (*right*) of Panic-TRAPed neurons in each neuronal cluster within and beyond the PHN.

Given that Panic-TRAPed neurons were labeled by tdTomato, we next analyzed the expression of tdTomato transcripts as an indicator of the neuronal ensemble activated during panic induction. Based on this approach, 2,495 Panic-TRAPed neurons were detected, covering multiple PHN neuronal clusters. For each neuronal cluster, we calculated the enrichment of Panic-TRAPed neurons by determining their proportion relative to the total number of cells in our dataset (Fig. 2n). Among the six PHN neuronal clusters, the N1 cluster (*Cbln2+*) contained the highest number and ratio of activated neurons; in contrast, the N5 (*Tac1*+) and N6 (*Sst*+) clusters contained the lowest number and ratio of activated neurons (Fig. 2o). These data suggest that N1 cluster (*Cbln2*+) in the PHN may be an important neuronal population involved in the panic-like defensive state in mice.

### Role of *Cbln2*+ PHN neurons in panic-like defensive state

Next, we examined the role of *Cbln2*+ PHN neurons (N1 cluster) in panic-like defensive state, using *Tac1*+ and *Sst*+ neurons (N5 and N6 clusters) as negative controls. Fluorescent *in situ* hybridization (FISH) confirmed the expression of *Cbln2* (Fig. 3a), *Sst* (Supplementary Fig. 4a) and *Tac1* (Supplementary Fig. 4d) in the PHN. The spatial distribution of neurons expressing these genes in the PHN was quantified (*Cbln2*: Fig. 3b; *Sst*: Supplementary Fig. 4b; *Tac1*: Supplementary Fig. 4e). About 4,000 *Cbln2*+ neurons were detected in the PHN, with a broad distribution in the rostral, intermediate, and caudal parts (Fig. 3b). At 2 h after panic induction, a considerable proportion (67% ± 7%, n = 5 mice) of *Cbln2+* neurons showed Fos expression, an indicator of neuronal activation (Fig. 3<u>, c & d</u>). Considering the high abundance of Fos-expressing *Cbln2*+ neurons, we hypothesized that *Cbln2+* PHN neurons may participate in the robot-induced panic-like state.

**Fig. 3.**
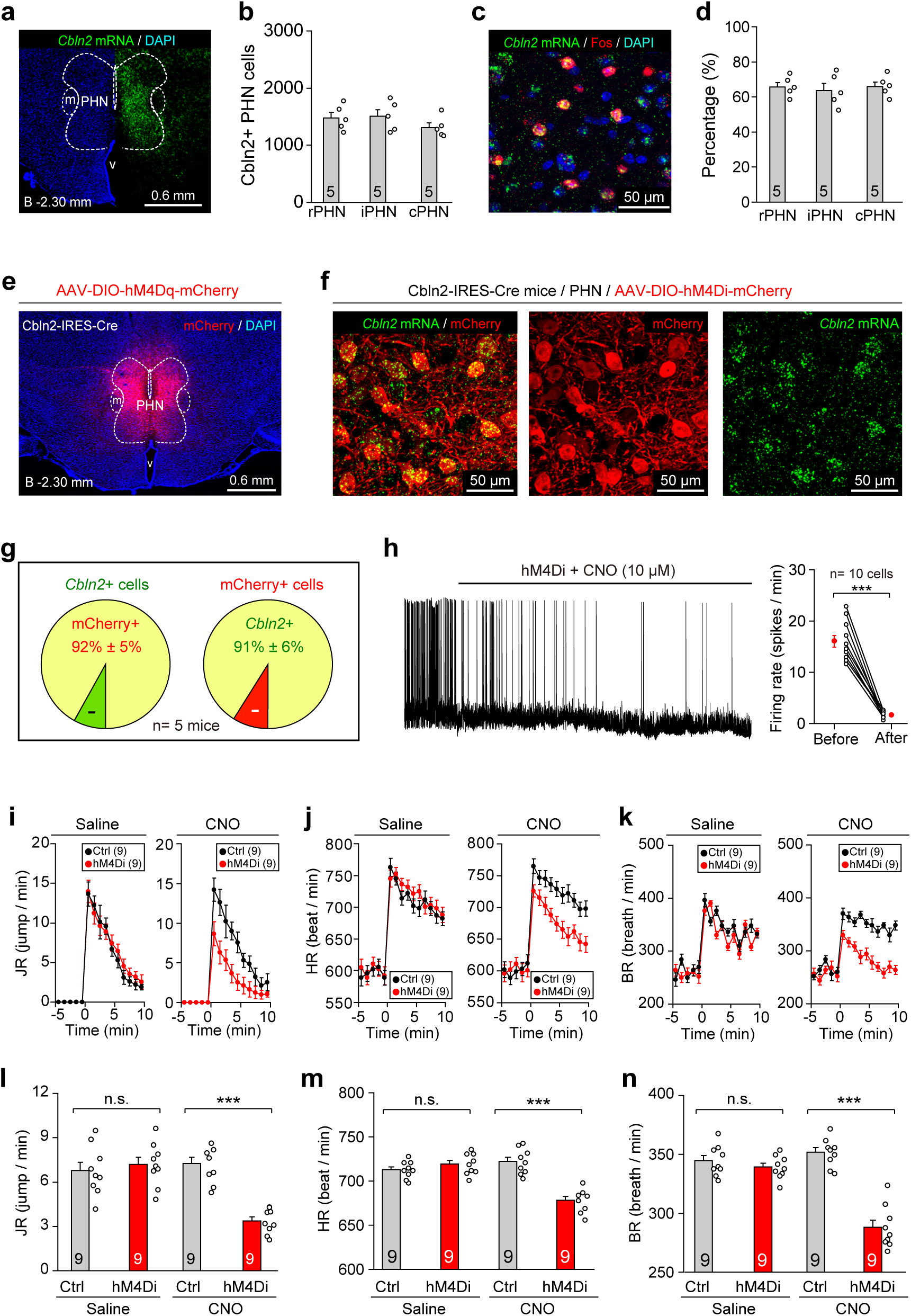
Identification of *Cbln2*+ PHN neurons as a key neuronal subtype for panic-like defensive responses. **(a,b)** Example coronal section (a) and quantitative analyses (b) showing distribution of *Cbln2*-expressing (*Cbln2*+) cells within PHN of WT mouse. **(c, d)** Example micrograph (c) and quantitative analyses (d) showing percentages of *Cbln2*+ PHN neurons expressing Fos after panic induction. **(e)** Example coronal brain section showing hM4Di-mCherry expression in PHN of Cbln2-IRES-Cre mice. **(f, g)** Example micrographs (f) and quantitative analyses (g) showing specificity and efficiency of Cbln2-IRES-Cre mice for labeling PHN cells expressing *Cbln2* mRNA. **(h)** Example trace of action potential firing (*left*) and quantitative analysis of firing rate (*right*) showing effectiveness of CNO at chemogenetically silencing hM4Di-expressing PHN neurons in acute brain slices. CNO was dissolved in ACSF (10 μM) and perfused in brain slices. **(i-k)** Time courses of jumping rate (i), heart rate (j), and breathing rate (k) before and during panic induction in mice treated with saline (control) or CNO (1 mg/kg) to chemogenetically inactivate *Cbln2*+ PHN neurons. **(l-n)** Quantitative analyses of jumping rate (l), heart rate (m), and breathing rate (n) during panic induction in mice treated with saline (control) or CNO (1 mg/kg) to chemogenetically inactivate *Cbln2*+ PHN neurons. Scale bars are labeled in graphs. Numbers of mice (b, d, g, i-n) and cells (h) are indicated in graphs. Data in (b, d, g-n) are means ± SEM. Statistical analyses (h, l-n) were performed using Student *t*-test (*** *P* < 0.001; n.s. *P* > 0.1). For *P*-values, see Supplementary Table 3.

To test this hypothesis, we examined whether *Cbln2*+ PHN neurons are required for the robot-induced panic-like state. AAV-DIO-hM4Di-mCherry (<u>Armbruster et al., 2007</u>) or AAV-DIO-mCherry (as control) was injected into the PHN of Cbln2-IRES-Cre mice (Fig. 3e). Specific expression of hM4Di-mCherry in *Cbln2*+ PHN neurons was validated (Fig. 3<u>, f & g</u>). The effectiveness of CNO in suppressing the firing of action potentials in *Cbln2*+ PHN neurons was confirmed by slice physiological recordings (Fig. 3h). We found that chemogenetic inactivation of *Cbln2*+ PHN neurons impaired robot-induced jumping escapes (Fig. 3<u>, i & l</u>), heart-rate increase (Fig. 3<u>, j & m</u>), and breathing-rate increase (Fig. 3<u>, k & n</u>). We also examined whether *Sst*+ and *Tac1*+ PHN neurons (N5 & N6 clusters) are required for the induction of panic-like state. AAV-DIO-hM4Di-mCherry was injected into the PHN of Sst-IRES-Cre and Tac1-IRES-Cre mice (<u>Taniguchi et al., 2011</u>; <u>Harris et al., 2014</u>) (Supplementary Fig. 4, c & f). Chemogenetic inactivation of *Sst*+ or *Tac1*+ PHN neurons did not significantly impair the robot-induced panic-like defensive state (*Sst*: Supplementary Fig. 4, g-l; *Tac1*: Supplementary Fig. 4, m-r). Thus, these findings suggest that *Cbln2*+ PHN neurons may be an important component in the central circuit module required for the panic-like defensive state in mice.

### Activation of *Cbln2*+ PHN neurons induces panic-like defensive state

We next explored whether *Cbln2*+ PHN neurons are causally linked to panic-like defensive state. First, we examined whether optogenetic activation of *Cbln2*+ PHN neurons evoked panic-associated behavioral and autonomic responses. We injected AAV-DIO-ChR2-mCherry (<u>Boyden et al., 2005</u>) in the PHN of Cbln2-IRES-Cre mice, followed by optical fiber implantation above the PHN (Fig. 4a). Specific expression of ChR2-mCherry in *Cbln2*+ PHN neurons was validated (Fig. 4<u>, b & c</u>). In acute brain slices, light pulses (473 nm, 5 mW, 2 ms, 10 or 20 Hz) reliably evoked action potentials in ChR2-expressing *Cbln2*+ PHN neurons (Fig. 4d). Optogenetic activation of *Cbln2*+ PHN neurons (473 nm, 5 mW, 20 Hz) induced jumping escapes (Fig. 4e; <u>Supplementary Video 5</u>), increased heart rate (Fig. 4f), and increased breathing rate (Fig. 4g). These light-evoked behavioral and autonomic responses depended on the frequency and laser power of light stimulation (Supplementary Fig. 5, a-c). Activation of *Cbln2*+ PHN neurons also induced real-time place aversion (Fig. 4h; Supplementary Fig. 5d). These observations suggest that brief optogenetic activation of *Cbln2*+ PHN neurons induces panic-like defensive state.

**Fig. 4.**
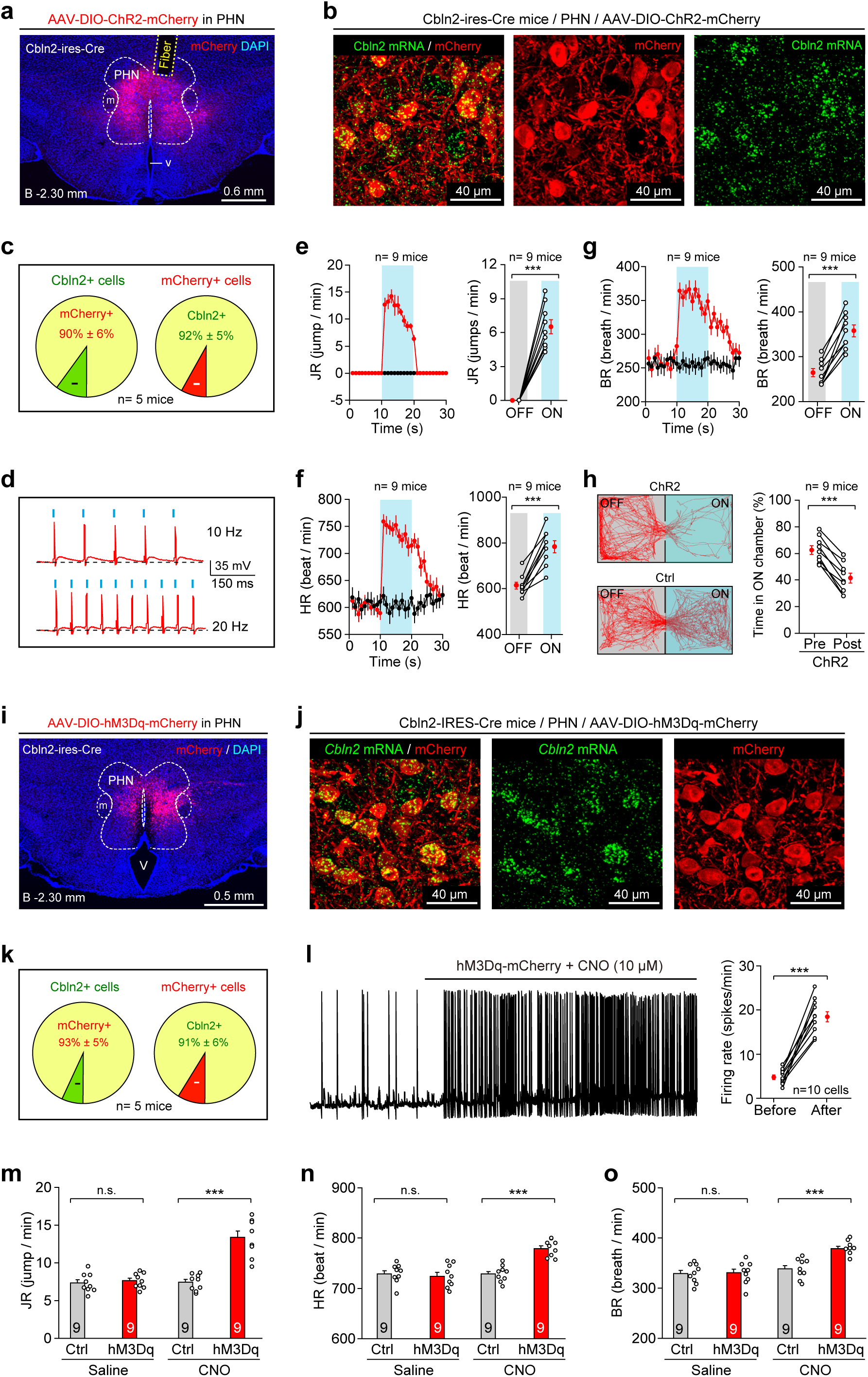
Activation of *Cbln2*+ PHN neurons evokes a panic-like defensive state. **(a)** Example coronal brain section showing optical fiber track above ChR2-mCherry+ PHN neurons in Cbln2-IRES-Cre mice. **(b, c)** Example micrographs (b) and quantitative analyses (c) showing specificity and efficiency of Cbln2-IRES-Cre mice for labeling PHN cells expressing *Cbln2* mRNA. **(d)** Light-pulse trains (473 nm, 2 ms, 5 mW, 10 Hz or 20 Hz) reliably evoked phase-locked spiking activity in ChR2-mCherry+ PHN neurons in acute brain slices. **(e-g)** Time courses (*left*) and quantitative analyses (*right*) of jumping rate (e), heart rate (f), and breathing rate (g) in mice with (ON) and without (OFF) light stimulation (20 ms, 5 mW, 20 Hz) of *Cbln2*+ PHN neurons. **(h)** Locomotion traces (*left*) and quantitative analyses (*right*) of real-time place aversion with (ON) and without (OFF) light stimulation of *Cbln2*+ PHN neurons. **(i)** Example coronal brain section showing hM3Dq-mCherry expression in PHN cells of Cbln2-IRES-Cre mice. **(j, k)** Example micrographs (j) and quantitative analyses (k) showing specificity and efficiency of Cbln2-IRES-Cre mice for labeling PHN cells expressing *Cbln2* mRNA. **(l)** Example trace of action potential firing (*left*) and quantitative analyses of firing rate (*right*) showing effectiveness of CNO to chemogenetically activate hM3Dq-expressing PHN neurons in acute brain slices. CNO was dissolved in ACSF (10 μM) and perfused in brain slices. **(m-o)** Quantitative analyses of jumping rate (m), heart rate (n), and breathing rate (o) during panic induction in mice with and without chemogenetic activation of *Cbln2*+ PHN neurons. Scale bars are labeled in graphs. Numbers of mice (e-h, m-o) or cells (l) are indicated in graphs. Data in (e-h, l-o) are means ± SEM. Statistical analyses (e-h, l-o) were performed using Student *t-*test (*** *P* < 0.001; n.s. *P* > 0.1). For *P*-values, see Supplementary Table 3.

Second, we examined whether chemogenetic activation of *Cbln2*+ PHN neurons promotes the robot-induced panic-like defensive state in mice. AAV-DIO-hM3Dq-mCherry was injected into the PHN of Cbln2-IRES-Cre mice, resulting in specific expression of hM3Dq-mCherry in *Cbln2*+ PHN neurons (Fig. 4<u>, i-k</u>). The effectiveness of CNO at evoking action potential firing in hM3Dq-mCherry+ neurons was validated using slice physiological recording (Fig. 4l). Strikingly, mice with chemogenetic activation of *Cbln2*+ PHN neurons exhibited more robot-induced jumping escapes (Fig. 4m; Supplementary Fig. 5e), as well as higher heart (Fig. 4n; Supplementary Fig. 5f) and breathing rates (Fig. 4o; Supplementary Fig. 5g) compared to control mice. However, chemogenetic activation of these neurons did not significantly increase the baseline levels of behavioral and autonomic responses (Supplementary Fig. 5, e-g). These results suggest that chemogenetic activation of *Cbln2*+ PHN neurons promotes the robot-induced defensive state in mice.

### Encoding of jumping escape vigor by *Cbln2*+ PHN neurons

To examine how *Cbln2*+ PHN neurons encode the panic-like defensive state, we injected AAV-DIO-jGCaMP7s into the PHN of Cbln2-IRES-Cre mice, resulting in specific expression of jGCaMP7s (<u>Dana et al., 2019</u>) in these neurons (Fig. 5<u>, a & b</u>). We monitored their GCaMP fluorescence using fiber photometry during panic induction in mice (Fig. 5c). We observed transient increases of GCaMP fluorescence (ΔF/F) that coincided with jumping escapes in individual test mouse (Fig. 5<u>, d & e</u>; <u>Supplementary Video 6</u>). In nine test mice (Fig. 5f), the increase in average GCaMP fluorescence of *Cbln2*+ PHN neurons (peak ΔF/F = 6.9% ± 0.9%, n = 9 mice) was tightly coupled to the initiation of jumping and reached a peak at 756 ± 62 ms after jumping. The increase in GCaMP fluorescence was not a motion artifact, as fluorescence in the EGFP-expressing *Cbln2*+ PHN neurons did not change during jumping escape (Supplementary Fig. 5, h & i).

**Fig. 5.**
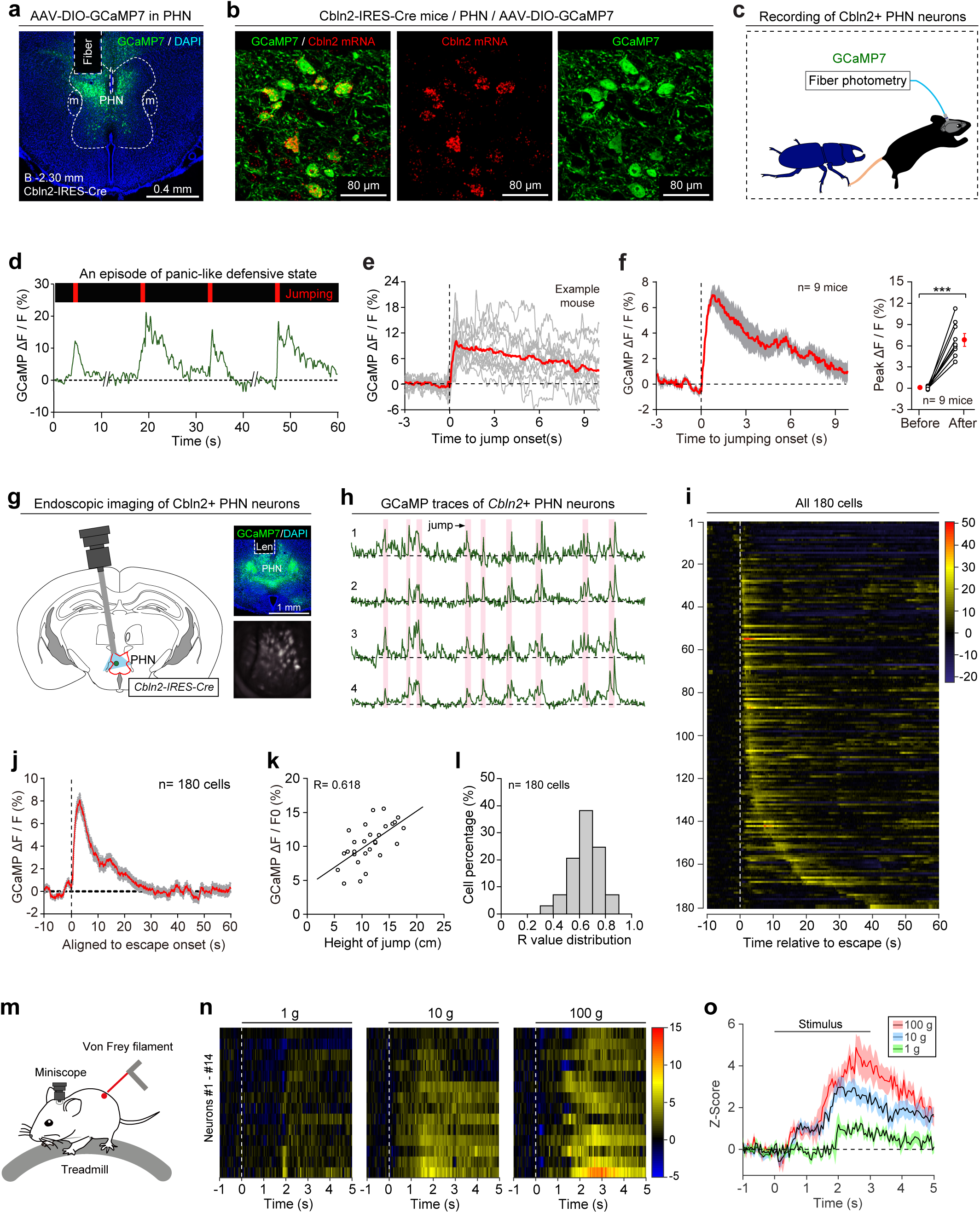
*Cbln2*+ PHN neurons encode jumping escape vigor. **(a)** Example coronal section showing optical fiber track above GCaMP7-expressing neurons in PHN of Cbln2-IRES-Cre mice. **(b)** Example micrographs from PHN showing specific expression of GCaMP7 in *Cbln2*+ PHN neurons. **(c)** Schematic diagram showing fiber photometry recordings of *Cbln2*+ PHN neurons in mice subjected to panic induction. **(d)** Normalized GCaMP fluorescence changes (green, ΔF/F) in *Cbln2*+ PHN neurons in parallel with jumping escape (red) in example mouse. Vertical red bars indicate jumping escape of mouse. **(e)** Individual traces (gray) and averaged trace (red) of normalized GCaMP fluorescence changes (ΔF/F) aligned with initiation of jumping in an example mouse. **(f)** Average GCaMP response curve (*left*) and quantitative analyses of peak GCaMP fluorescence changes (*right*) of all tested mice (n= 9). **(g)** Schematic diagram (*left*) and example micrographs (*right*) showing GRIN lens implantation and imaging of *Cbln2*+ PHN neurons with the endoscope system. **(h)** Example traces of normalized GCaMP fluorescence changes in individual cells during panic induction in an example mouse. Red shaded areas indicate occurrence of jumping escape. **(i, j)** Heat-map (i) and average traces (j) of GCaMP responses of all 180 *Cbln2*+ PHN neurons in seven mice aligned with initiation of jumping escape. **(k)** Scatter plot showing GCaMP responses (ΔF/F) of an example *Cbln2*+ PHN neuron linearly correlated with jumping escape height. Value of correlation coefficient for this example cell was 0.618. **(l)** Distribution of correlation coefficients of all 180 cells. **(m)** Schematic diagram showing endoscope imaging of individual *Cbln2*+ PHN neurons in response to application of von Frey filaments to the back of test mice. **(n)** Heat-map peri-stimulus time histogram (PSTH) of Z-score GCaMP fluorescence changes in individual *Cbln2*+ PHN neurons to mechanical stimuli applied with von Frey filaments (1, 10, and 100 g). **(o)** Average PSTH of Z-score GCaMP fluorescence changes in all *Cbln2*+ PHN neurons to mechanical force applied with different intensities on back. Scale bars are labeled in graphs. Numbers of mice (f) and cells (i, j, n) are indicated in graphs. Data in (f, j, o) are means ± SEM. Statistical analyses (f) were performed using Student t-test (*** *P* < 0.001). For *P*-values, see Supplementary Table 3.

To examine whether single *Cbln2*+ PHN neurons respond to jumping escape, *in vivo* Ca^2+^ imaging of individual *Cbln2*+ PHN neurons was performed in freely moving mice with a head-mounted miniature endoscope and an implanted gradient index (GRIN) lens (<u>Ghosh et al., 2011</u>) (Fig. 5g). When the mice displayed jumping escape, most *Cbln2*+ PHN neurons (180/202) exhibited an increase in GCaMP signals relative to baseline (Fig. 5, h and i; <u>Supplementary Video 7</u>). When the GCaMP signals of all recorded *Cbln2*+ PHN neurons were averaged, the peak GCaMP signal reached approximately 8% after jumping escape (Fig. 5j). For each neuron, the amplitude of the GCaMP signal during individual trials showed a linear correlation with jumping escape height in mice (Fig. 5k). The correlation coefficients (R value) of 180 cells were normally distributed, with an average value of 0.657 (Fig. 5l). These findings suggest that the activity of *Cbln2*+ PHN neurons may encode jumping escape vigor.

Considering that the height of jumping escape may reflect the mechanical strength exerted during collision, we examined the response of *Cbln2*+ PHN neurons to mechanical force. The GCaMP responses of individual *Cbln2*+ PHN neurons were recorded in head-fixed mice on a treadmill, with mechanical stimuli applied by poking the back of the mice with von Frey filaments (Fig. 5m). We found that *Cbln2*+ PHN neurons (n= 14 neurons) responded to mechanical force with different intensities (1g, 10g, 100g) (Fig. 5n). Quantitative analyses indicated that these neurons exhibited graded responses, showing a preference for stronger mechanical force (Fig. 5o). Together, these results suggest that *Cbln2*+ PHN neurons respond to mechanical stimuli generated by mouse-robot collisions during panic induction.

### Synaptic inputs of *Cbln2*+ PHN neurons to regulate panic-like defensive state

To reveal the origin of mechanical signals in *Cbln2*+ PHN neurons, retrograde tracing with recombinant G-deleted rabies virus (RV) (<u>Wickersham et al., 2007</u>) was performed to map the circuits that monosynaptically innervate *Cbln2*+ PHN neurons (Fig. 6<u>, a & b</u>; Supplementary Fig. 6, a-c). We found that *Cbln2*+ PHN neurons were monosynaptically innervated by the parabrachial nucleus (PBN), lateral septum (LS), dorsal raphe nucleus (DRN), and other brain areas (Fig. 6c; Supplementary Fig. 6, d & e). The PBN receives neural signals of mechanical pain from the dorsal horn of the spinal cord (<u>Huang et al., 2019</u>; <u>Chiang et al., 2020</u>; <u>Choi et al., 2020</u>). Furthermore, LS projections to the hypothalamus are implicated in the regulation of anxiety-like behaviors in mice (<u>Anthony et al., 2014</u>; <u>Wang et al., 2023</u>). These studies suggest that PBN-PHN and LS-PHN circuits may regulate the robot-induced panic-like defensive state.

**Fig. 6.**
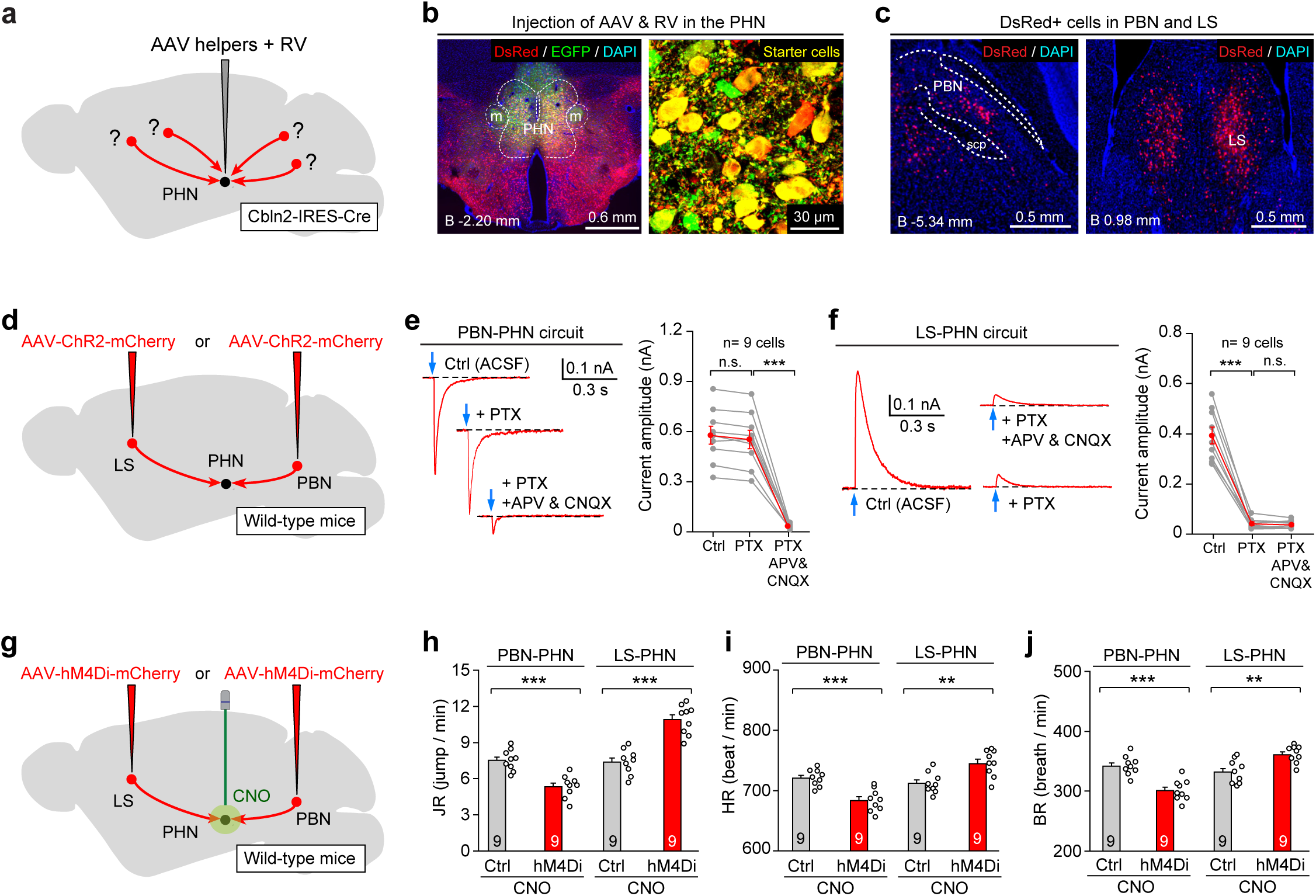
Inputs of *Cbln2*+ PHN neurons for regulating panic-like defensive state. **(a)** Schematic diagram showing injection of AAV-helpers and RV to retrogradely label neurons monosynaptically projecting to *Cbln2*+ PHN neurons. For detailed information on AAV-helpers and RV, see Supplementary Fig. 5a. **(b)** Example coronal brain section (*left*) and magnified area (*right*) showing starter cells in PHN of Cbln2-IRES-Cre mice. For single-channel micrographs of starter cells, see Supplementary Fig. 5c **(c)** Example micrographs showing retrogradely labeled DsRed+ neurons in PBN (*left*) and LS (*right*). For DsRed+ cells in other areas and quantitative data, see Supplementary Fig. 5d and 5e. **(d)** Schematic diagram showing injection of AAV-ChR2-mCherry in LS or PBN of WT mice. **(e, f)** Example traces (*left*) and quantitative analyses (*right*) showing effects of picrotoxin (PTX, 50 μM) and APV (50 μM)/CNQX (20 μM) on postsynaptic currents recorded from PHN neurons in acute slices in response to photostimulation (2 ms, 5 mW) of PBN-PHN (e) or LS-PHN (f) circuits. **(g)** Schematic diagram showing strategy to chemogenetically inactivate PBN-PHN or LS-PHN pathways. **(h-j)** Quantitative analyses of jumping rate (h), heart rate (i), and breathing rate (j) during panic induction in mice with CNO infusion to chemogenetically inactivate PBN-PHN and LS-PHN circuits. Scale bars are labeled in graphs. Numbers of mice (h-j) or cells (e, f) are indicated in graphs. Data in (e, f, h-j) are means ± SEM. Statistical analyses (e, f, h-j) were performed using Student *t*-test (*** *P* < 0.001; n.s. *P* > 0.1). For *P*-values, see Supplementary Table 3.

To test this possibility, we characterized the synaptic properties of PBN-PHN and LS-PHN circuits. AAV-ChR2-mCherry was injected into the PBN or LS of wild-type (WT) mice (Fig. 6d). Light-evoked postsynaptic currents (PSCs) from PHN neurons were then recorded in acute brain slices containing the PHN. Notably, PSCs in PHN neurons (n= 9) evoked by light stimulation (473 nm, 2 ms, 5 mW) of the PBN-PHN circuit were selectively blocked by glutamate receptor antagonists APV (50 μM) and CNQX (20 μM), suggesting that the PBN-PHN circuit was primarily glutamatergic (Fig. 6e). In contrast, PSCs in PHN neurons (n= 9) evoked by light stimulation (473 nm, 2 ms, 5 mW) of the LS-PHN circuit were selectively eliminated by the GABAa receptor antagonist picrotoxin (50 μM), suggesting that the LS-PHN circuit was predominantly GABAergic (Fig. 6f).

Next, we examined the functional significance of PBN-PHN and LS-PHN circuits in panic-like defensive state. AAV-hM4Di-mCherry was bilaterally injected into the PBN or LS of WT mice, followed by implantation of a pair of cannulas above the PHN (Fig. 6g). Infusion of CNO (1 mM, 500 nl, 10 min) to chemogenetically inactivate the PBN-PHN circuit significantly impaired the robot-induced panic-like defensive state; in contrast, chemogenetic inactivation of the LS-PHN circuit significantly promoted the panic-like defensive state (Fig. 6<u>, h-j</u>; Supplementary Fig. 6, f-h). These findings suggest that the PBN-PHN and LS-PHN circuits differentially regulate the robot-induced panic-like defensive state in mice.

### *Cbln2*+ PHN-PAG pathway to regulate panic-like defensive state

We next mapped the axonal projections of *Cbln2*+ PHN neurons by injecting AAV-DIO-EGFP into the PHN of Cbln2-IRES-Cre mice (Fig. 7a). The EGFP-expressing *Cbln2*+ PHN neurons (Fig. 7b) projected to the PAG in the midbrain, Barrington’s nucleus (Bar) / locus coeruleus (LC) in the pons, and other brain regions (Fig. 7c; Supplementary Fig. 7, a & b). PAG neurons are involved in defensive behaviors (<u>Tovote et al., 2016</u>). The Bar is a pontine micturition center for urination (<u>Yao et al., 2018</u>). The LC is implicated in the regulation of arousal state (<u>Carter et al., 2010</u>). These studies suggest that the *Cbln2*+ PHN-PAG and PHN-Bar/LC circuits may be involved in the robot-induced panic-like defensive state.

**Fig. 7.**
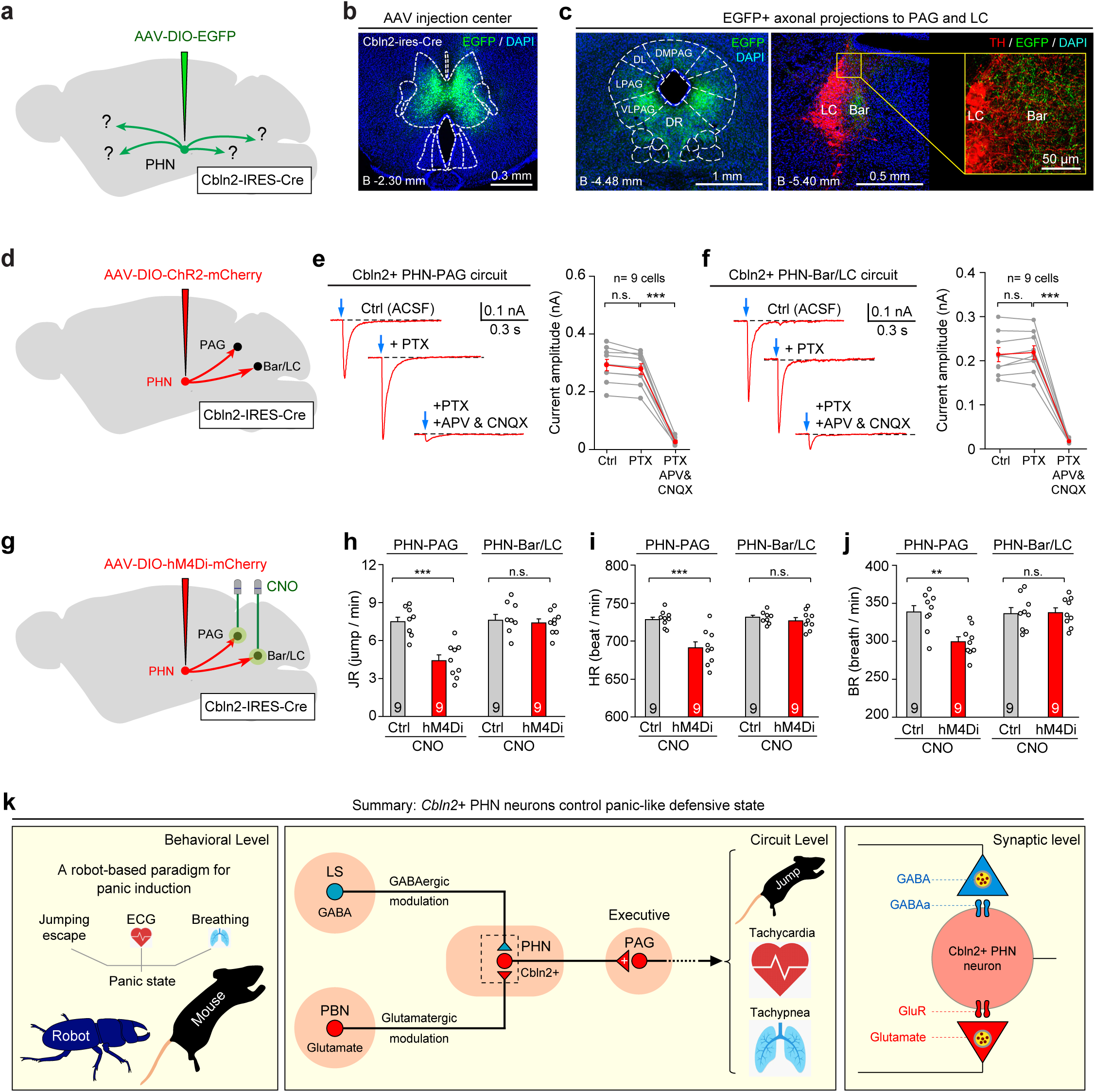
Outputs of *Cbln2*+ PHN neurons for regulating panic-like defensive state. **(a)** Schematic diagram showing injection of AAV-DIO-EGFP to anterogradely label projections of *Cbln2*+ PHN neurons to different brain areas. **(b)** Example coronal brain section showing expression of EGFP in PHN of Cbln2-IRES-Cre mice. **(c)** Example micrographs showing EGFP+ axons in PAG (*left*) and Bar/LC (*right*). For projections in other brain areas and quantitative data, see Supplementary Fig. 6a and 6b. **(d)** Schematic diagram showing injection of AAV-DIO-ChR2-mCherry into PHN of Cbln2-IRES-Cre mice. **(e, f)** Example traces (*left*) and quantitative analyses (*right*) showing effects of picrotoxin (PTX, 50 μM) and APV (50 μM)/CNQX (20 μM) on postsynaptic currents evoked by photostimulation (2 ms, 5 mW) of *Cbln2*+ PHN-PAG (e) or PHN-Bar/LC (f) circuits. **(g)** Schematic diagram showing strategy to chemogenetically inactivate *Cbln2*+ PHN-PAG or PHN-Bar/LC pathway. **(h-j)** Quantitative analysis of jumping rate (h), heart rate (i), and breathing rate (j) during panic induction in mice with CNO infusion to chemogenetically inactivate *Cbln2*+ PHN-PAG and PHN-Bar/LC pathways. **(k)** Summary graphs. *Left*, a robot-based paradigm for panic induction. *Middle*, A molecularly defined circuit module, composed of *Cbln2*+ PHN neurons and their input/output circuits, for regulating panic-like defensive state in mice. *Right*, synaptic integration in *Cbln2*+ PHN neurons. Scale bars are labeled in graphs. Numbers of mice (h-j) or cells (e, f) are indicated in graphs. Data in (e, f, h-j) are means ± SEM. Statistical analyses (e, f, h-j) were performed using Student *t*-test (*** *P* < 0.001; n.s. *P* > 0.1). For *P*-values, see Supplementary Table 3.

To test this possibility, we characterized the synaptic properties of these two downstream circuits. AAV-DIO-ChR2-mCherry was injected into the PHN of Cbln2-IRES-Cre mice (Fig. 7d). Light-evoked PSCs from PAG or Bar/LC neurons were then recorded in acute brain slices containing the PAG or Bar/LC. We found that light-evoked PSCs from both PAG (n= 9 cells) and Bar/LC (n= 9 cells) neurons were selectively blocked by glutamate receptor antagonists APV/CNQX but not by the GABAa receptor antagonist picrotoxin (Fig. 7<u>, e & f</u>), suggesting that both circuits were primarily glutamatergic.

We then explored how the *Cbln2*+ PHN-PAG and PHN-Bar/LC circuits regulate the panic-like defensive state. AAV-DIO-hM4Di-mCherry was bilaterally injected into the PHN of Cbln2-IRES-Cre mice, followed by implantation of a pair of cannulas above the PAG or Bar/LC (Fig 7g; Supplementary Fig. 7c). Infusion of CNO (1 mM, 500 nl, 10 min) to chemogenetically inactivate the *Cbln2*+ PHN-PAG circuit significantly impaired the robot-induced panic-like state; in contrast, chemogenetic inactivation of the *Cbln2*+ PHN-Bar/LC circuit did not significantly impair the panic-like state (Fig. 7<u>, h-j</u>; Supplementary Fig. 7, d-f). Overall, these findings suggest that the *Cbln2*+ PHN-PAG pathway, rather than the PHN-Bar/LC pathway, may play a primary role in robot-induced behavioral and autonomic control of panic-like state.

## DISCUSSION

Panic is a strong defensive state with important physiological functions and pathological relevance. The synaptic and circuit mechanisms underlying panicogenesis remain poorly understood. An appropriate experimental paradigm is critical for addressing this question. Here, we have developed a robot-based paradigm for panic induction, and thereby revealed a molecularly defined circuit module that integrates diverse panic-associated neurochemical inputs and drives panic-like defensive state in mice (Fig. 7n).

### A robot-based paradigm to induce panic-like defensive state in mice

As described by the classical “predatory imminence continuum” theory, defensive modes in prey are organized as a function of physical distance between prey and predator (<u>Fanselow and Lester, 1988</u>). In mammals, the three defensive modes (pre-encounter, post-encounter, and circa-strike) are well mapped onto states of anxiety, fear, and panic (<u>McNaughton and Corr, 2004</u>; <u>Mobbs et al., 2007</u>; <u>Hoffman et al., 2022</u>). Specifically, when prey is physically attacked by a predator, it may exhibit a “circa-strike” response, characterized by either fight or flight, depending on escape availability. Accordingly, we established a robot-based paradigm by placing a test mouse and an electrically powered robot (a predator surrogate) together in the same arena. The size of the arena (15 × 15 cm) was small enough for the robot to frequently collide with the mouse, thus inducing circa-strike jumping escapes (flight) (<u>Supplementary Video 1</u>). Quantitative analyses indicated that the test mice exhibited repetitive jumping escapes in parallel with sustained increases in heart and breathing rates. These behavioral and physiological responses in mice are analogous to those observed in human during panic attacks, such as tachycardia, tachypnea, and strong desire to escape (<u>American Psychiatric Association, 2013</u>). Importantly, these responses were reduced by chronic treatment with fluoxetine, a selective serotonin reuptake inhibitor for treating panic disorder (<u>Quagliato et al., 2018</u>). Thus, we defined the robot-induced behavioral and physiological responses as a panic-like defensive state in mice.

Our collision-based panicogenic paradigm offers several advantages over other methods for inducing panic. First, it holds higher ethological relevance, as the panic-like defensive state relies on actual and imminent collisions between the robot and mouse (Fig. 1). In ethology, actual and imminent collisions symbolize danger for both human and animals (<u>Billington et al., 2010</u>; <u>Yilmaz and Meister 2013</u>). Second, the use of mice as the animal model in this paradigm facilitates the application of molecular tools for analyzing the circuit and synaptic mechanisms underlying panic attacks (<u>Luo et al., 2018</u>).

### A critical role of *Cbln2*+ PHN neurons in panic-like defensive state

In human subjects, deep brain stimulation in the posterior hypothalamus elicited panic attacks, suggesting a possible involvement of this region in panic induction (<u>Schoenen et al., 2005</u>; <u>Bartsch et al., 2008</u>). In consistent with these human studies, we found that the PHN in mice is critical for panic induction (Fig. 1). Using snRNA-seq and functional analyses, we identified *Cbln2*+ PHN neurons as a key neuronal population involved in the regulation of the panic-like defensive state in mice (Fig. 2 & 3). In rats, the PHN was known to be involved in stress-induced hypothalamic-pituitary-adrenal responses (<u>Nyhuis et al., 2016</u>) and tachycardia evoked by corticotropin releasing factor (<u>Gao et al., 2016</u>). Together, the observations in different species (human, mice, and rats) suggest that the PHN may be an evolutionarily conserved brain area involved in panicogenesis.

However, our results do not rule out the possible involvement of other brain areas in the regulation of panic-like defensive state. Previous studies have revealed multiple brain areas that participate in the regulation of fear (<u>Gross and Canteras, 2012</u>), including the ventromedial hypothalamic nucleus (<u>Wang et al., 2015</u>; <u>Kunwar et al., 2015</u>), dorsal premammillary nucleus (<u>Wang et al., 2021</u>), dorsomedial hypothalamic nucleus (<u>Johnson and Shekhar, 2006</u>), amygdala (<u>Ziemann et al., 2009</u>), and bed nucleus of stria terminalis (<u>Taugher et al., 2014</u>). The key neuronal populations in these brain areas that regulate panic state warrants further study.

### Afferents of *Cbln2*+ PHN neurons to regulate panic-like defensive state

In this study, we explored the regulatory roles of the PBN-PHN and LS-PHN pathways in the panic-like defensive state. The PBN is known to receive neural signals related to mechanical pain from the dorsal horn of the spinal cord (<u>Huang et al., 2019</u>; <u>Chiang et al., 2020</u>; <u>Choi et al., 2020</u>). Our results suggest that the PBN-PHN pathway may transmit collision-associated mechanical stimuli to the PHN to regulate the panic-like defensive state. These findings are consistent with recent research demonstrating that *Adcyap1*+ PBN neurons are involved in regulating the panic-like state (<u>Kang et al., 2024</u>). Further research is needed to elucidate the potential role of *Adcyap1*+ PBN neurons in the robot-induced panic-like defensive state.

The LS projections to the hypothalamus are implicated in the regulation of anxiety-like behaviors in mice (<u>Anthony et al., 2014</u>; <u>Wang et al., 2023</u>). In this study, we found that inhibition of the GABAergic LS-PHN pathway significantly increased the robot-induced panic-like defensive state, supporting the hypothesis that the GABA system may play an important regulatory role in panic attacks (<u>Zwanzger and Rupprecht, 2005</u>; <u>Johnson et al., 2014</u>).

In addition to the PBN and LS, *Cbln2*+ PHN neurons also received monosynaptic inputs from the ventral subiculum and other brain areas (Supplementary Fig. 5d). Recent research has revealed that a neural circuit extending from the ventral subiculum to the anterior hypothalamic nucleus plays an essential role in anxiety-like behavioral avoidance (<u>Yan et al., 2022</u>). Exploring the potential influence of the ventral subiculum on the panic-like defensive state via its monosynaptic inputs to *Cbln2*+ PHN neurons would be a valuable area of research.

### Role of PAG and Bar/LC in panic-like defensive state

Our findings indicated that the *Cbln2*+ PHN-PAG pathway is essential for inducing panic, while the *Cbln2*+ PHN-Bar/LC pathway appears to be dispensable in this context.

This aligns with previous human studies. For example, deep brain stimulation of the PAG in human evoked panic attack (<u>Nashold et al., 1969</u>). In contrast, stimulation of the LC in human results in NE release and promotion of alertness, without generation of anxiety, fear, or panic (<u>Libet and Gleason, 1994</u>). This suggests that the *Cbln2*+ PHN-PAG pathway may be the primary circuit for regulating the panic state, whereas the PHN-Bar/LC pathway may regulate non-cardiorespiratory responses, such as arousal (<u>Carter et al., 2010</u>; <u>Schwarz and Luo, 2015</u>) and urination (<u>Yao et al., 2018</u>) in response to threats.

### Limitations of the study

Our study has a few limitations. <u>First</u>, we did not examine the role of *Cbln2*+ PHN neurons in panic-like states induced by interoceptive stimuli, such as carbon dioxide or sodium lactate, which requires further exploration in future study. Considering that PBN neurons may serve as a central hub to detect danger from both external and internal environments (<u>Palmiter, 2018</u>), it will be important to test whether PBN-PHN pathway may also be involved in mediating panicogenesis induced by CO2 or sodium lactate. <u>Second</u>, we did not examine the possibility of sexual dimorphism in PHN neurons in panic. As panic disorder is known to exhibit sex differences (<u>Sheikh et al., 2002</u>), it would be interesting to examine whether the role of PHN neurons in panic induction is sex-dependent in mice.

## ACKNOWLEDGEMENTS

This work was supported by the National Natural Science Foundation of China (31925019 to PC) and National Key R&D Program of China (2021ZD0202701 to P.C.). We declare no conflicts of interest. All data are archived in NIBS.

## AUTHOR CONTRIBUTIONS

Conceptualization, P.C., Q.W., D.L., Z.X., F.Z., and Y.T.; Investigation, M.Z., L.Z., Z.C., S.Z., X.C., M.H., X.L., H.G., X.G., D.G., Y.L., and Z.X.; Writing – original draft, P.C. and Q.W.; Funding acquisition, P.C. and Q.W.; Resources, H.M. and D.L.; Methodology, D.L.; Supervision, P.C., D.L, F.Z., and Q.W.

## METHODS

### Animals

All experimental procedures were conducted following protocols approved by the Administrative Panel on Laboratory Animal Care (NIBS2022M0028) in the National Institute of Biological Sciences (NIBS) in Beijing, China. The various mouse lines, including Tac1-IRES-Cre (<u>Harris et al., 2014</u>), Sst-IRES-Cre (<u>Taniguchi et al., 2011</u>), Rosa26-iDTR (<u>Buch et al., 2005</u>), Mrgprd-CreERT2 (<u>Olson et al., 2017</u>), Ai3 (<u>Madisen et al., 2010</u>), Ai9 (<u>Madisen et al., 2010</u>), and FosTRAP2 mice (<u>Allen et al., 2017</u>) were imported from the Jackson Laboratory (JAX Mice and Services). Cbln2-IRES-Cre mice were generated by our own lab and was characterized previously (<u>Xie et al., 2021</u>). Mice were maintained on a circadian 12-h light/12-h dark cycle with food and water available *ad libitum*. Mice were housed in groups (3–5 animals per cage) before separation three days prior to virus injection. After virus injection, each mouse was individually housed in a cage for at least 3 weeks before subsequent experiments. To avoid potential sex-specific differences, we used male mice only.

### Stereotaxic injection of viral vectors

#### Viral vectors

Adeno-associated virus (AAV) serotype (AAV2/9) was used. The AAVs and RV used in the present study are listed in Supplementary Table 6. Viral particles were purchased from Shanghai Taitool Bioscience Inc. and Brain VTA Inc. The viral vector titers before dilution were in the range of 0.8–1.5 × 10^13^ viral particles/ml. The final titer used for AAV injection was 5 × 10^12^ viral particles/ml.

#### Stereotaxic injection

Stereotaxic injection was performed according to our published work (<u>Xie et al., 2022</u>). Mice were anesthetized with an intraperitoneal injection of tribromoethanol (125–250 mg/kg). Standard surgery was performed to expose the brain surface above the PHN, PBN, LS, PAG, LC, and DRN. Coordinates used for PHN injection were: bregma −2.30 mm, lateral ±0.25 mm, and dura −4.50 mm. Coordinates used for PBN injection were: bregma −5.34 mm, lateral ±1.30 mm, and dura −2.25 mm. Coordinates used for LS injection were: bregma +0.50 mm, lateral ±0.25 mm, and dura −2.50 mm. Coordinates used for PAG injection were: bregma −4.50 mm, lateral ±0.30 mm, and dura −1.50 mm. Coordinates used for LC injection were: bregma −5.40 mm, lateral ±0.75 mm, and dura −2.50 mm. Coordinates used for DRN injection were: bregma −4.50 mm, lateral ±0.15 mm, and dura −2.25 mm. The AAVs were injected with a glass pipette connected to a Nano-liter Injector 201 (World Precision Instruments, Inc.) at a slow flow rate of 0.15 μl/min to avoid potential damage to local brain tissue. The pipette was withdrawn at least 20 min after viral injection. For experimental designs, see Supplementary Table 1.

### FosTRAP experiments

#### Using FosTRAP2::Ai9 mice to label panic-associated neurons in whole brain

4-hydroxy-tamoxifen (4-OHT) was dissolved in corn oil (20 mg/ml) overnight on a shaker at 37 °C. FosTRAP2::Ai9 mice were intraperitoneally injected with 4-OHT at a dose of 30 mg/kg. At 1 h after injection, the mice were placed in the arena and subjected to the paradigm for panic induction for 30 minutes (Panic-TRAP). The control mice were placed in the arena for 30 minutes without exposure to the robot (Ctrl-TRAP). 14 days after TRAP experiment, the mice were perfused for histological analyses of tdTomato-labeled panic-associated neurons in the brain.

#### Using FosTRAP2 mice to inactivate panic-associated PHN neurons

AAV-DIO-hM4Di-mCherry or AAV-DIO-mCherry (control) was bilaterally injected into the PHN of *FosTRAP2* mice on Day 1. On Day 7, 4-OHT was intraperitoneally injected into the *FosTRAP2* mice at a dose of 30 mg/kg. At 1 h after injection, the mice were placed in the arena and subjected to the paradigm for panic induction for 30 minutes. On Day 28 (3 weeks after Panic-TRAP), the mice were subjected to panic induction again, followed by perfusion 2 h after induction. Coronal sections (40 μm) containing the hypothalamic brain areas were collected for immunohistochemical analysis of mCherry and Fos.

### Single-nucleus RNA sequencing

#### Single-nucleus isolation and snRNA-seq

To isolate single nuclei from PHN of Fos-TRAP2::Ai9 mice treated with panic attacks, we immersed the tissues in ice-cold homogenization buffer containing 250 mM sucrose, 25 mM KCl, 5 mM MgCl_2_, 10 mM Tris-HCl, 0.1 mM DTT, 1% BSA, 0.1% NP-40, and ribonuclease inhibitor (0.4 U/μl). The tissues were then mechanically homogenized using a glass homogenizer for 10 cycles to obtain a single-nucleus suspension for snRNA-seq. Single-nucleus RNA-seq libraries were prepared using DNBSEQ technology platforms (BGI Genomics, China) and the DNBelab C4 Single-Cell Library Prep Set (MGI Tech, #1000021082). Briefly, single-nucleus suspensions were introduced into a microfluidic device for droplet generation. Following emulsion breakage and bead collection, reverse transcription and cDNA amplification were performed to generate barcoded libraries. The cDNA products were then sheared to short fragments with a length of 250 to 400 bp, and indexed sequencing libraries were constructed according to the manufacturer’s protocol. The quality of sequencing libraries was assessed using the Qubit ssDNA Assay Kit (Thermo Fisher Scientific, #Q10212). DNBs were loaded into patterned nano arrays and sequenced on the ultrahigh-throughput DIPSEQ T-series sequencer.

#### Sequencing data preprocessing

Raw sequencing reads from DIPSEQ T-series sequencer were filtered and demultiplexed using PISA (v0.10) (https://github.com/shiquan/PISA). To identify the positive cells in Fos-TRAP2::Ai9 mice, we inserted the tdTomato sequences (download from : <u>tdTomato Sequence and Map (snapgene.com)</u>) into GRCm38 reference genome. Reads were aligned to the modified GRCm38 mouse genome using STAR (v2.7.4a) (https://github.com/alexdobin/STAR) and sorted using Sambamba (v0.7.0) (<u>Tarasov et al., 2015</u>). A nucleus versus gene UMI count matrix was generated by PISA for downstream analysis. Cells expressing tdTomato mRNA were judged to be positive cells.

#### Quality control and data integration

To ensure the quality of the data, we applied the following criteria to select cells for further analysis: (1) the percentage of mitochondrial gene counts was less than 10%; (2) the cells had more than 1000 UMIs and between 200 and 7,000 detected genes; (3) we used DoubletFinder (v2.0.2) (<u>McGinnis et al., 2019</u>) to remove doublets. DoubletFinder creates artificial doublets by averaging the transcriptional profiles of randomly chosen cell pairs. It then predicted which cells were likely to be doublets based on their proximity to the artificial doublets in the gene expression space. The neighborhood size (pK) was determined using the mean-variance normalized bimodality coefficient (BCMVN) for each sample. The number of artificial doublets (pN) was set to 0.25.

We integrated the multiple datasets using the fast integration of reciprocal principal component analysis (RPCA) in Seurat (v4.0.1) (<u>Hao et al., 2021</u>). First, we identified the top 2000 variable features in each dataset and performed principal components analysis (PCA). We then used the “FindIntegrationAnchors” function to identify potential anchors between the datasets. For each pair of datasets, we used “IntegrateData” function to correct batch effects and integrate datasets.

#### Dimensionality reduction and clustering

To reduce the dimensionality of the integrated datasets, we performed PCA and applied the first 50 principal components (PCs) to calculate uniform manifold approximation and projection (UMAP). To identify cell clusters, we used a shared nearest neighbor (SNN)-based clustering algorithm. This algorithm calculates the K-nearest neighbors (K=20) for each cell, constructs an SNN graph, and optimizes the modularity function to determine clusters. We adopted a stepwise clustering strategy for each dataset. First, we clustered the cells with the Louvain algorithm using “FindClusters” function at a resolution of 0.5 to distinguish neurons from non-neuronal cells. Next, we subsetted neurons from diverse single-cell datasets for further reclustering. To identify cell type-specific genes, we compared the neuronal clusters pairwise. We assigned cell identities by cross-referencing their marker genes with known neuronal subtype markers.

#### Differential expression analysis

To identify differentially expressed genes (DEGs) between cell types, we performed pairwise comparisons of cell groups using the “FindAllMarkers” function in Seurat. The Wilcoxon rank-sum test was used to calculate log_2_ fold changes and *P* values for each gene, which were corrected using the Bonferroni method. Only genes with an average log_2_-transformed difference greater than 0.25, a *P* value less than 0.05 were considered as DEGs.

### Genetic ablation of *Mrgprd*+ neurons

The ablation of *Mrgprd*+ neurons was conducted according to our previous work (<u>Xie et al., 2022</u>). To ablate *Mrgprd*+ neurons, a two-step strategy of drug injections was used. First, tamoxifen was injected intraperitoneally for eight consecutive days at a dosage of 75 mg/kg in 14-day-old Mrgprd-CreERT2/iDTR/Ai3 male mice. Tamoxifen injection at this dosage at this developmental stage was shown to induce Cre expression in Mrgprd-expressing DRG neurons with high specificity (88.1% ± 1%) and high efficiency (92.9% ± 4.6%) (<u>Olson et al., 2017</u>). Second, 3 weeks after the last dose of tamoxifen injection, diphtheria toxin or saline (as a control) was injected intraperitoneally for three consecutive days at a dosage of 4 mg/kg in the same group of mice. The panic-like defensive responses was measured 3 weeks after the last dose of diphtheria toxin injection. The efficiency of genetic ablation of Mrgprd+ neurons was tested by immunostaining of EYFP in the dorsal root ganglia (DRG) at the lumbar and sacral segments.

### Preparation of behavioral tests

After AAV injection, the mice were housed individually for 3 weeks before the behavioral tests and handled daily by the experimenters for at least three days. On the day of the behavioral test, the mice were transferred to the testing room and were habituated to the room conditions for 3 h before the experiments started. The apparatus was cleaned with 20% ethanol to eliminate odor cues from other mice. All behavioral tests were conducted during the same circadian period (13:00–19:00). All behaviors were scored by researchers who were blind to the animal treatments. The illuminance in the “light” condition for behavior was 50 lux, whereas the ‘dark’ condition was ∼0.002 lux, as measured with an illuminance meter (RTR Optoelectronics Technology).

### Measuring panic-like defensive responses

#### Robot for panic induction

The robot used for panic induction was HEXBUG Scarab (4 x 6 x 8 cm, 23 g), a high-speed (30 cm/s) beetle-like robotic bug that skitters around on six angled legs. In each experiment, new batteries were loaded to the robot for optimal induction of panic state in mice.

#### Measuring jumping escapes

To measure jumping escapes, the arena (15 cm x 15 cm) was placed on top of a force plate. The force-plate actometer device used in this study was designed according to a published work (<u>Fowler et al., 2001</u>). The mouse was allowed to freely explore the arena for 10 minutes before the robot was placed in the arena. The robot moved in random directions and frequently collided with the mouse, resulting in its circa-strike jumping escapes. The vertical force generated by each jumping escape was recorded by the force plate, with a sampling rate at 100 Hz. For each jumping escape, the trace showed two high-amplitude peaks that corresponded to jump up and falling down, respectively (Fig. 1b). To verify the jumping escapes recorded by force plate, mouse behavior was also recorded by a high-speed camera. The timing and number of jumping escapes were analyzed offline by using a custom MATLAB script for data analysis.

#### Measuring heart-beat rate

The mice were implanted with DSI implantable probe (HD-X11) according to the manual provided by the manufacturer (DSI Data Science). After implantation, the mice were treated with antibiotics and allowed to recover in home cage for 5 days. On the day of ECG recording, the mice were placed in the arena (15 cm x 15 cm) for free exploration for 30 minutes. The experiment was started when heart rate of mice returned to the resting state (550∼600 bpm). After placing the robot in the arena, the moving robot randomly collided with the test mouse and caused jumping escapes. The two receivers (RPC-1) was vertically positioned in parallel with the walls of the arena to receive ECG signal. The ECG signal was fed into the Data exchange Matrix (DEM), transmitted to the computer, and recorded with New Ponemah Acquisitions Software. The heart rate was measured offline with Spike2 Software (CED Inc.).

#### Measuring breathing rate

The method to record breathing followed the method in our published work (<u>Liu et al., 2017</u>). One week before respiration recording, mice under anesthesia were implanted with an intranasal cannula (inner diameter 0.5 mm, outer diameter 0.7 mm). The cannula was fixed to the skull with dental cement and was protected with the canula cap. On the day of recording, the intranasal cannula was connected to a pressure transducer (MPX5050, Motorola Semiconductors H.K. Ltd) via a piece of polyethylene tube. During the experiment, the intranasal respiratory pressure in freely moving mice was converted to voltage signal by the pressure transducer in real time. The voltage signal was amplified 100×, low pass filtered at 100 Hz, and digitized at 8000 Hz using a data acquisition unit (Micro 1401, CED Inc.). The rate of breathing was analyzed offline with Spike2 Software (CED Inc.).

### Chemogenetic inactivation/activation

#### AAV injection and CNO preparation

For chemogenetic inactivation/activation of PHN neurons, AAV-DIO-hM4Di-mCherry (for inactivation), AAV-DIO-hM3Dq-mCherry (for activation), or AAV-DIO-mCherry (as control) was bilaterally injected into the PHN of individual Cre mouse lines. Chemogenetic experiment was performed 3 weeks after viral injection. Clozapine N-oxide (CNO) stock solution was prepared in saline at 5 mg/ml. To prepare working solution for experiments, CNO was diluted 1:10 with saline (0.5 mg/ml).

#### Effects of chemogenetic inactivation on panic-like defensive responses

Mice with neurons expressing hM4Di-mCherry or mCherry (control) were intraperitoneally treated with CNO (1 mg/kg) and were allowed for rest in home-cage for 20 minutes. The mice were placed in an arena (15 cm x 15 cm) and allowed for free exploration for 10 minutes. The robot was then placed in the arena, causing collisions with the test mouse. The behavior of mouse was recorded with a high-speed camera. The jumping escape was measured by the force-plate actometer. The ECG was recorded by DSI telemetry system. The breathing was recorded with the respiration recording system.

#### Effects of chemogenetic activation on panic-like defensive responses

Mice with neurons expressing hM3Dq-mCherry or mCherry (control) were intraperitoneally treated with CNO (1 mg/kg) and were allowed for rest for 20 minutes in home-cage. The mice were placed in an arena (15 cm x 15 cm) and allowed for free exploration for 10 minutes. The robot was placed in the arena, resulting in collisions with the test mouse. The behavior of mouse was recorded with a high-speed camera. The jumping escape was measured by the force-plate actometer. The ECG was recorded by DSI telemetry system. The respiration was recorded by the respiration recording system.

#### Cannula implantation and CNO infusion

For chemogenetic inactivation of PBN-PHN or LS-PHN circuit, AAV-hSyn-hM4Di-mCherry was injected in the PBN or LS, followed by implantation of a pair of cannulae above the PHN. For chemogenetic inactivation of Cbln2+ PHN-PAG or PHN-Bar/LC circuit, AAV-hSyn-DIO-hM4Di-mCherry was injected in the PHN, followed by implantation of a pair of cannulae above the PAG or Bar/LC of Cbln2-IRES-Cre mice. The inner and outer diameters of the cannula were 150 μm and 300 μm, respectively. The cannula was fixed to the skull using acrylic cement. During drug infusion, the cannula was connected with a catheter filled with CNO (10 mM) or saline for injection. The other end of the catheter was connected to a Hamilton syringe controlled by an infusion pump to drive the delivery of CNO (500 nl) or saline (control).

#### Effects of chemogenetic inactivation on panic-like defensive responses

After infusion, mice were allowed to rest in home-cage for 20 minutes. The mice were then placed in an arena (15 cm x 15 cm) and allowed for freely exploring the arena for 10 minutes. Then the robot was placed in the arena, resulting in collisions with the test mouse. The behavior of mouse was recorded with a high-speed camera. The jumping escape was measured by the force-plate actometer. The heart-beat rate was measured by telemetry DSI system. The breathing rate was measured by the respiration recording system. At the end of the experimental session, the test mice were perfused, and the coronal brain sections containing PHN, PAG or Bar/LC were inspected for the presence of cannula track. The mice with no cannula track above the targeted brain areas were rejected from further analysis.

### Optogenetic activation

#### AAV injection and optical fiber Implantation

AAV-hSyn-DIO-ChR2-mCherry was unilaterally injected into the PHN of Cbln2-IRES-Cre mice. At 30 min after AAV injection, a ceramic ferrule with an optical fiber (200 µm in diameter, N.A. 0.37) was implanted with the optical fiber tip on top of the PHN (bregma −2.30 mm, lateral ±0.25 mm, dura −4.50 mm). The ferrule was secured on the skull with dental cement. After optical fiber implantation, the skin was sutured, and antibiotics were applied to the surgical wound.

#### Measuring panic-like defensive responses induced by optogenetic activation

Three weeks after AAV injection, panic-like defensive responses to photostimulation of Cbln2+ PHN neurons were measured. The test mouse with optical fiber was allowed to freely explore the arena for 10 minutes. Then light stimulation (473 nm, 10 s, 20 ms) was applied, with variable light power (1 and 5 mW) and frequency (2 and 20 Hz). The jumping escape was recorded by the force-plate actometer. The ECG of mice was recorded by DSI telemetry system. The breathing was recorded by implanted nasal tube connected to the transducer. In each experiment, three trials per mouse per condition were used to reduce the variability of experimental results.

#### Measuring photostimulation-induced real-time place aversion

The test mouse with optical fiber was initially placed in the two-chamber arena for 10 minutes and the time spent in each chamber was measured. After 24 hours, the mouse with optical fiber was placed in the two-chamber arena for 10 minutes again. Light stimulation (20 Hz, 5 mW, 20 ms, 2 s) was automatically triggered when the mouse entered its preferred chamber. The locomotion of the test mouse in the two-chamber arena was recorded with a camera above the arena. The time spent in each chamber was measured.

### Fiber photometry

#### AAV injection and optical fiber implantation

AAV-hSyn-DIO-GCaMP7s was injected into the PHN of Cbln2-IRES-Cre mice. At 30 min after AAV injection, a ceramic ferrule with an optical fiber (200 µm in diameter, N.A. 0.37) was implanted with the optical fiber tip on top of the PHN (bregma −2.30 mm, lateral ±0.25 mm, dura −4.50 mm). The ferrule was secured on the skull with dental cement. After optical fiber implantation, the skin was sutured, and antibiotics were applied to the surgical wound.

#### Fiber photometry system

A fiber photometry system (Thinker Tech, Nanjing, China) was used for recording fluorescence signals of GCaMP from Cbln2+ PHN neurons (<u>Gunaydin et al., 2014</u>). To induce fluorescence signals, a laser beam from a laser tube (488 nm) was reflected by a dichroic mirror, focused by a 10× lens (N.A. 0.3), and coupled to an optical commutator. A 2-m optical fiber (230-μm diameter, N.A. 0.37) guided the light between the commutator and implanted optical fiber. To minimize photo bleaching, the power intensity at the fiber tip was adjusted to 0.02 mW. The fluorescence of GCaMP7s (<u>Dana et al., 2019</u>) was band-pass filtered (MF525-39, Thorlabs) and collected using a photomultiplier tube (R3896, Hamamatsu). An amplifier (C7319, Hamamatsu) was used to convert the photomultiplier tube current output to voltage signals, which were further filtered through a low-pass filter (40 Hz cut-off; Brownlee 440). The analogue voltage signals were digitalized at 100 Hz and recorded by a Power 1401 digitizer and Spike2 software (CED, Cambridge, UK).

#### Fiber photometry recording

Three weeks after optical fiber implantation, fiber photometry was performed to record fluorescence signal of GCaMP from Cbln2+ PHN neurons in freely moving mice. The behavior of mice was monitored by a high-speed camera. A flashing LED triggered by a 1-s square-wave pulse was simultaneously recorded to synchronize the video and fluorescence signals. The test mice were subjected to robot-induced panic induction. After the experiments, the optical fiber tip sites above the PHN were histologically examined in each mouse.

### GRIN lens implantation and miniscope Ca^2+^ imaging

#### AAV injection and GRIN lens implantation

To monitor the activities of *Cbln2*+ PHN neurons before and during panic induction, microendoscopic imaging was conducted according to our previous study (<u>Hou et al., 2023</u>). AAV-hsyn-DIO-jGCaMP7s was injected into the PHN of Cbln2-IRES-Cre mice. Two weeks after viral injection, a second surgery was conducted to expose the brain surface above PHN. A GRIN lens (0.5 mm in diameter, 6.1 mm in length, Inscopix) was held by a GRIN lens holder (Inscopix) and was slowly (100 μm/min) inserted to the target brain area. To avoid damaging the superior sagittal sinus, the brain tissue above the PHN was not aspirated. To better ascertain the correct depth of GRIN lens, the lens was connected to the imaging system of miniscope to monitor the GCaMP fluorescence. The GCaMP fluorescence gradually increased and reached the optimal intensity, indicating the appropriate depth of lens. Then the lens was anchored to the skull with dental cement. The silicon adhesive (Kwik-Sil, WPI) was used to cap the lens to avoid damage.

#### Baseplate attachment

Two weeks after GRIN lens implantation, a miniature microscope (nVista, Inscopix) and a baseplate (1050-002192, Inscopix) were attached on top of the lens. The height of the microscope relative to the lens was adjusted until an in-focus imaging field was obtained. The baseplate was secured to the skull with dental cement. A baseplate cover (1050-002193, Inscopix) was implemented to protect the lens from dust or damage. After surgery, mice were individually housed for recovery. Before microendoscopic Ca^2+^ imaging, the test mice were adapted to the microscope for 3 days, with half hour each day.

#### Acquisition of Ca^2+^ imaging data

Microendoscopic Ca^2+^ imaging was conducted by using a miniaturized fluorescence microscopy unit linked to the data acquisition device (nVista, Inscopix). Before the imaging sessions, the microscope was attached to the baseplate and secured with locking screw. The behavior of mice were monitored by a side-view camera. A data acquisition device with Inscopix Data Acquisition Software (ver. 1.3.1) was used to establish a connection between the microscope, host computer and camera. Synchronization between mouse behavior and Ca^2+^ imaging was implemented by using the external transistor-transistor logic (TTL) signals from the data acquisition device. The images (1440 × 1080 pixels) were acquired with a fluorescence power of 0.375-1.0 mW/mm^2^ (473 nm) at a 20 Hz frame rate, and the pixel size was approximately 0.625 μm.

#### Recording of Ca^2+^ imaging during panic induction

On the day of panic induction, the test mouse was equipped with the miniscope and placed in the arena. After ten minutes, the test mouse was subjected to robot-based panic induction. The jumping escapes and microendoscopic Ca^2+^ imaging were simultaneously recorded. To minimize photobleaching, the recording period was 10 mins for each session. The GCaMP signals of single neurons were measured by normalizing GCaMP fluorescence (ΔF/F).

#### Recording Ca^2+^ imaging before and during mechanical stimuli

Head-fixed awake mice with the GRIN lens connected to the miniscope system were allowed to stand on a circular treadmill. Then, noxious mechanical stimuli were applied by the von Frey filaments to the back of mouse body while microendoscopic Ca^2+^ imaging were simultaneously recorded. The GCaMP signals of single neurons were measured by normalizing GCaMP fluorescence (ΔF/F) and aligned with the initiation of mechanical stimuli.

#### Data analysis

Imaging data were processed by the Inscopix data processing software (Version 1.6.0). Raw images were then spatially down-sampled by a binning factor of 2, followed by motion correction. Ca^2+^ signal traces of Cbln2+ PHN neurons were extracted with the extended constrained non-negative matrix factorization (CNMF-E) in MATLAB (MathWorks). All traces were denoised and deconvoluted using the integrated CNMF-E program. Every extracted cell was manually checked for circular spatial footprints, and Ca^2+^ transients were characterized by sharp rises and slow decays. We then excluded certain regions of interests from the analysis based on anatomy (size, shape, or vicinity to the edge of lens), low signal-to-noise ratio, or large overlap in signal and spatial location with other neurons (>60% spatial overlap, and Pearson’s correlation coefficient > 0.6 between the traces across the entire session). The Z-scores were computed as (F(t) − Fm)/s.d., where F(t) is the averaged fluorescence signal (ΔF) value at time t, and Fm and s.d. are the mean and standard deviation of the ΔF values over a baseline period of –2 s to 0 s. All traces from the same experimental condition were aligned and sorted based on the average Ca^2+^ fluorescence intensity difference of all the neurons. The mean activity trace was calculated by averaging across the traces of all neurons, with at least three trials for each stimulus type.

### Cell-type-specific RV tracing

A modified RV-based three-virus system was used for mapping whole-brain inputs to Cbln2+ PHN neurons (<u>Wickersham et al., 2007</u>). All viruses, including AAV2/9-CAG-DIO-tdTomato-2A-TVA (5 × 10^12^ viral particles per ml), AAV2/9-CAG-DIO-RVG (5 × 10^12^ viral particles/ml), and EnvA-pseudotyped, glycoprotein (RG)-deleted, and EGFP-expressing RVs (RV-EvnA-ΔG-EGFP) (5.0 × 10^8^ viral particles/ml), were packaged and provided by BrainVTA Inc. (Wuhan, China). A mixture of AAV2/9-CAG-DIO-tdTomato-2A-TVA and AAV2/9-CAG-DIO-RVG (1:1, 200 nl) was injected into the PHN of Cbln2-IRES-Cre mice bilaterally. Two weeks after AAV helper injection, RV-EvnA-ΔG-EGFP (300 nl) was injected into the same location in the PHN of Cbln2-IRES-Cre mice in a biosafety level-2 lab facility. Starter neurons were characterized by the co-expression of tdTomato and EGFP, which were restricted in the PHN. One week after injection of RV, the mice were perfused with saline, followed by 4% paraformaldehyde (PFA) in phosphate-buffered saline (PBS). The brain was post-fixed in 4% PFA for 8 h and then incubated in PBS containing 30% sucrose until they sank to the bottom. Coronal brain sections (40-μm thickness) were prepared using a cryostat (Leica CM1900). All sections were collected and stained with 4’,6-diamidino-2-phenylindole (DAPI). The sections were imaged with an Olympus VS120 epifluorescence microscope (10× objective lens) and analyzed with ImageJ software. For quantification of subregions, boundaries were based on the Allen Institute’s reference atlas. We selectively analyzed retrogradely labeled dense areas. The fractional distribution of total cells labeled with RV was measured.

### Cell-type-specific anterograde tracing

For cell-type-specific anterograde tracing of *Cbln2*+ PHN neurons, AAV-DIO-EGFP (200 nl) was injected into the PHN of Cbln2-IRES-Cre mice. The mice were then maintained in a cage individually. Three weeks after viral injection, the mice were perfused with saline, followed by 4% PFA in PBS. The brains were post-fixed in 4% PFA for 8 h and then incubated in PBS containing 30% sucrose until they sank to the bottom. Coronal brain sections (40-μm thickness) were prepared using a cryostat (Leica CM1900). All coronal sections were collected and stained with primary antibodies against EGFP and DAPI. The coronal brain sections were imaged with an Olympus VS120 epifluorescence microscope (10× objective lens).

### Slice physiological recording

Preparation of acute brain slices was performed according to previous study (<u>Liu et al., 2017</u>). Brain slices containing the PHN were prepared from adult mice anesthetized with isoflurane before decapitation. Brains were rapidly removed and placed in ice-cold oxygenated (95% O_2_ and 5% CO_2_) cutting solution (228 mM sucrose, 11 mM glucose, 26 mM NaHCO_3_, 1 mM NaH_2_P O_4_, 2.5 mM KCl, 7 mM MgSO_4_, and 0.5 mM CaCl_2_). Coronal brain slices (400-μm thickness) were cut using a vibratome (VT 1200S, Leica Microsystems, Wetzlar, Germany). The slices were incubated at 28 °C in oxygenated ACSF (125 mM NaCl, 2.5 mM KCl, 1.25 mM NaH_2_PO_4_, 1.0 mM MgCl_2_, 25 mM NaHCO_3_, 15 mM glucose, and 2.0 mM CaCl_2_) for 30 min (∼305 mOsm, pH 7.4). The slices were then kept at room temperature under the same conditions for 30 min before transfer to the recording chamber at room temperature. The ACSF was perfused at 1 ml/min. The acute brain slices were visualized with a 40× Olympus water immersion lens, differential interference contrast optics (Olympus Inc., Japan), and a CCD camera.

Patch pipettes were pulled from borosilicate glass capillary tubes (Cat #64-0793, Warner Instruments, Hamden, CT, USA) using a PC-10 pipette puller (Narishige Inc., Tokyo, Japan). For recording of action potentials (current clamp), pipettes were filled with solution (135 mM K-methanesulfonate, 10 mM HEPES, 1 mM EGTA, 1 mM Na-GTP, 4 mM Mg-ATP, pH 7.4). For recording of postsynaptic currents (voltage clamp), pipettes were filled with solution (135 mM CsCl, 10 mM HEPES, 1 mM EGTA, 2 mM QX-314 chloride, 1 mM Na-GTP, 4 mM Mg-ATP, pH 7.4). The resistance of pipettes varied between 3.0–3.5 MΩ. The current and voltage signals were recorded with MultiClamp 700B and Clampex 10 data acquisition software (Molecular Devices). After establishment of the whole-cell configuration and equilibration of the intracellular pipette solution with the cytoplasm, series resistance was compensated to 10–15 MΩ. Recordings with series resistances > 15 MΩ were rejected. An optical fiber (200-μm diameter) was used to deliver light pulses, with the fiber tip positioned 500 μm above the brain slices. Laser power was adjusted to 5 mW before each experiment. Pulse onset, duration, and frequency of light stimulation were controlled by a programmable pulse generator attached to the laser system. Light-evoked postsynaptic currents from ChR2-mCherry-negative neurons in the PHN were triggered by a single light-pulse (2 ms) in the presence of 4-aminopyridine (4-AP, 20 μM) and tetrodotoxin (TTX, 1 μM). APV (50 μM), CNQX (20 μM), picrotoxin (PTX, 50 μM) were perfused with ACSF to examine the neurotransmitter receptor type of optically evoked postsynaptic currents. 5-HT (10 μM) and WAY100635 (1 μM) were perfused with ACSF to examine the 5-HT receptor type to mediate 5-HT effects on action potential firing.

### Immunohistochemical analysis

Mice were anesthetized with isoflurane and sequentially perfused with saline and PBS with 4% PFA. Brains were incubated in PBS containing 30% sucrose until they sank to the bottom. Cryostat sections (40 μm) were collected, incubated overnight with blocking solution (PBS containing 10% goat serum and 0.7% Triton X-100), then treated with primary antibodies diluted with blocking solution for 3–4 h at room temperature. Primary antibodies used for immunohistochemical analysis are displayed in Supplementary Table 6. Primary antibodies were washed three times with washing buffer (PBS containing 0.7% Triton X-100) before incubation with secondary antibodies (tagged with Cy2, Cy3, or Cy5; dilution 1:500; Life Technologies Inc.) for 1 h at room temperature. Sections were then washed three times with washing buffer, stained with DAPI, washed with PBS, transferred onto Super Frost slides, and mounted under glass coverslips with mounting media.

Sections were imaged with an Olympus VS120 epifluorescence microscope (10× objective lens) or a Zeiss LSM 710 confocal microscope (20× and 60× oil-immersion objective lens). Samples were excited by 488-, 543-, or 633-nm lasers in sequential acquisition mode to avoid signal leakage. Saturation was avoided by monitoring pixel intensity under Hi-Lo mode. Confocal images were analyzed with ImageJ software.

### RNA *in situ* hybridization

The method for RNA in situ hybridization followed our published work (<u>Chen et al., 2020</u>). Mice were perfused with 0.1% DEPC-(Sigma, D5758) treated PBS, followed by DEPC-treated PBS containing 4% PFA (PBS-PFA). Brains were post-fixed in DEPC-treated PBS-PFA solution overnight and then placed in DEPC-treated 30% sucrose solution at 4 °C for 30 h. Brain sections (30-μm thickness) were prepared using a cryostat (Leica, CM3050S) and collected in DEPC-treated PBS. Fluorescence *in situ* hybridization (FISH) was performed as described previously (<u>Chen et al., 2020</u>), with minor modifications. Briefly, brain sections were rinsed with DEPC-treated PBS, permeabilized with DEPC-treated 0.1% Tween 20 solution (in PBS) and DEPC-treated 2 × SSC containing 0.5% Triton. Brain sections were then treated with H_2_O_2_ solution and acetic anhydride solution to reduce nonspecific FISH signals. After 2 h of incubation in prehybridization buffer (50% formamide, 5 × SSC, 0.1% Tween20, 0.1% CHAPS, and 5 mM EDTA in DEPC-treated water) at 65 °C, the brain sections were hybridized with hybridization solution containing mouse anti-sense cRNA probes (digoxigenin labeling) for Tac1, Cck, and Npy at 65 °C for 20 h. The cDNA primer sequences for the cRNA probes were the same as those in the ISH data from the Allen Brain Atlas (https://mouse.brain-map.org/), as listed in Supplementary Table 6. After washing, the brain sections were incubated with anti-digoxigenin-POD, Fab fragments (1:400, Roche, 11207733910) at 4 °C for 30 h, and FISH signals were detected using a TSA Plus Cyanine 3 Kit (NEL744001KT, PerkinElmer). To visualize tdTomato and mCherry signals, the brain sections were incubated with primary antibodies against mCherry at 4 °C for 24 h and then with Alexa Fluor® 488-conjugated goat anti-rabbit secondary antibodies (1:500, A11034, Invitrogen) at room temperature for 2 h. The brain sections were mounted and imaged using a Zeiss LSM780 confocal microscope or the Olympus VS120 Slide Scanning System.

### Cell-counting strategies in brain areas and quantitative analyses

The cell-counting strategies are summarized in Supplementary Table 2. For counting cells in the PHN, we collected 40-μm coronal sections from bregma −1.8 mm to −2.7 mm for each mouse. Five evenly spaced (200 μm) sections were sampled for immunohistochemistry or FISH to label cells positive for different markers. We acquired fluorescent images (10× objective, Olympus FV1200 microscope) of the PHN, followed by cell counting with ImageJ software.

## QUANTIFICATION AND STATISTICAL ANALYSIS

All experiments were performed with anonymized samples in which the experimenter was unaware of the experimental conditions of the mice. For statistical analyses of experimental data, Student *t*-test were used. The “n” used in these analyses represents number of mice or cells. Detailed information on statistical analyses is provided in the figure legends and Supplementary Table 3.

## RESOURCE AND DATA AVAILABILITY

Further information and requests for resources and reagents should be directed to and will be fulfilled by the Lead Contact, Dr. Peng Cao. The Cbln2-IRES-Cre knock-in mouse driver line generated in this study is free to share with other investigators upon request. Requests for raw data and original codes reported in this paper will be shared by the lead contact upon request. The accession number for the snRNA-seq data reported in this paper is GEO: GSE252356. Additional information on snRNA-seq data will be shared by Dr. Qingfeng Wu upon request.

## LEGENDS

**Supplementary Fig. 1.**
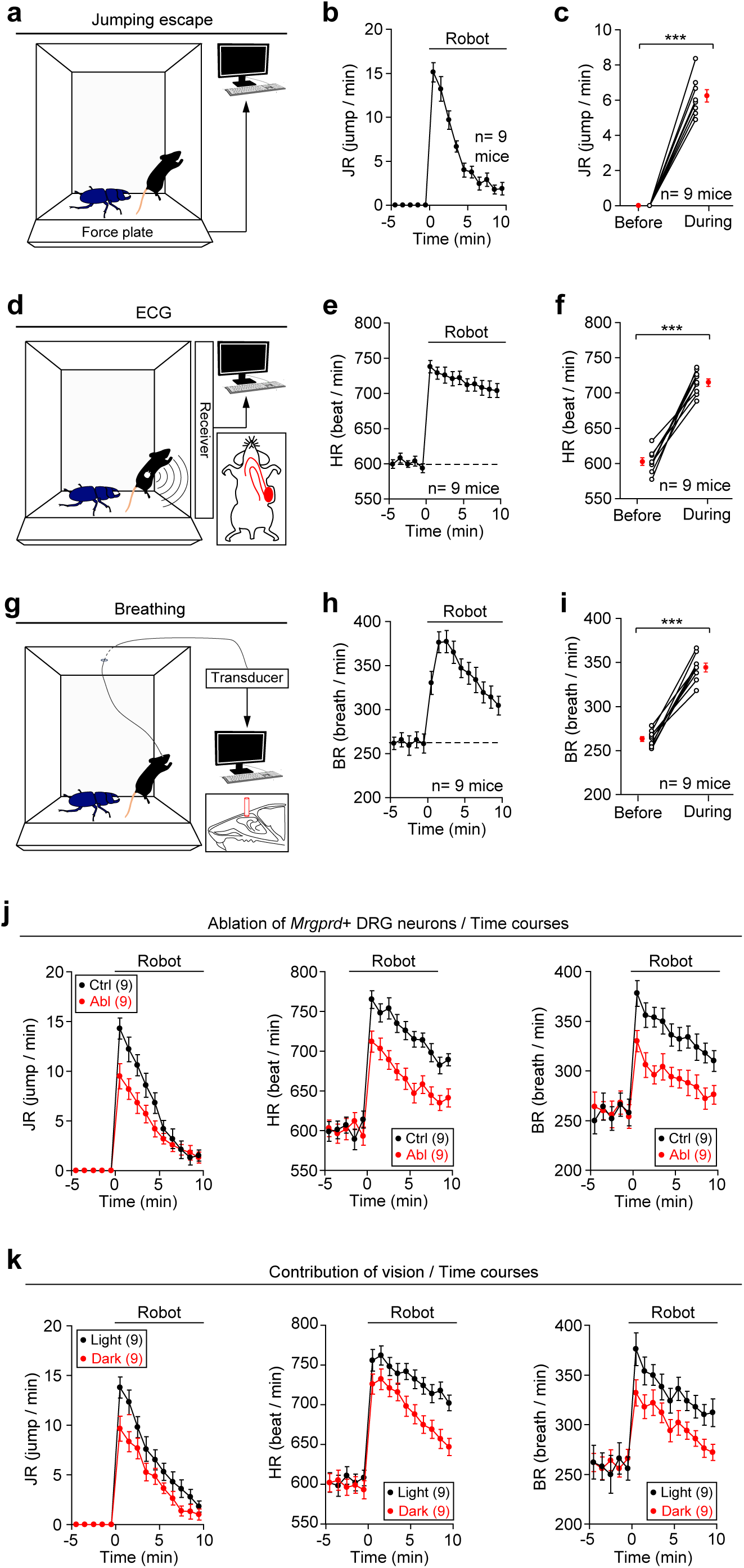
Experimental paradigm to induce panic-like defensive state in mice. **(a)** Schematic diagram showing recording of jumping escape using force-plate actometer. **(b, c)** Time course (b) and quantitative analyses (c) of jumping rate in mice before and during panic induction. **(d)** Schematic diagram showing recording of ECG using DSI telemetry system. **(e, f)** Time course (e) and quantitative analyses (f) of heart rate in mice before and during panic induction. **(g)** Schematic diagram showing recording of breathing using a cannulae implanted above the nasal cavity. **(h, i)** Time course (h) and quantitative analyses (i) of breathing rate in mice before and during panic induction. **(j)** Time courses of jumping rate (*left*), heart rate (*middle*), and breathing rate (*right*) in mice with and without ablation of *Mrgprd*+ DRG neurons before and during panic induction. **(k)** Time courses of jumping rate (*left*), heart rate (*middle*), and breathing rate (*right*) in mice before and during panic induction in an arena with (50 lux) and without (0.002 lux) light illumination. Numbers of mice (b, c, e, f, h, i, j, k) are indicated in graphs. Data in (b, c, e, f, h, i, j, k) are means ± SEM. Statistical analyses (c, f, i) were performed using Student *t*-test (*** *P* < 0.001). For *P*-values, see Supplementary Table 3.

**Supplementary Fig. 2.**
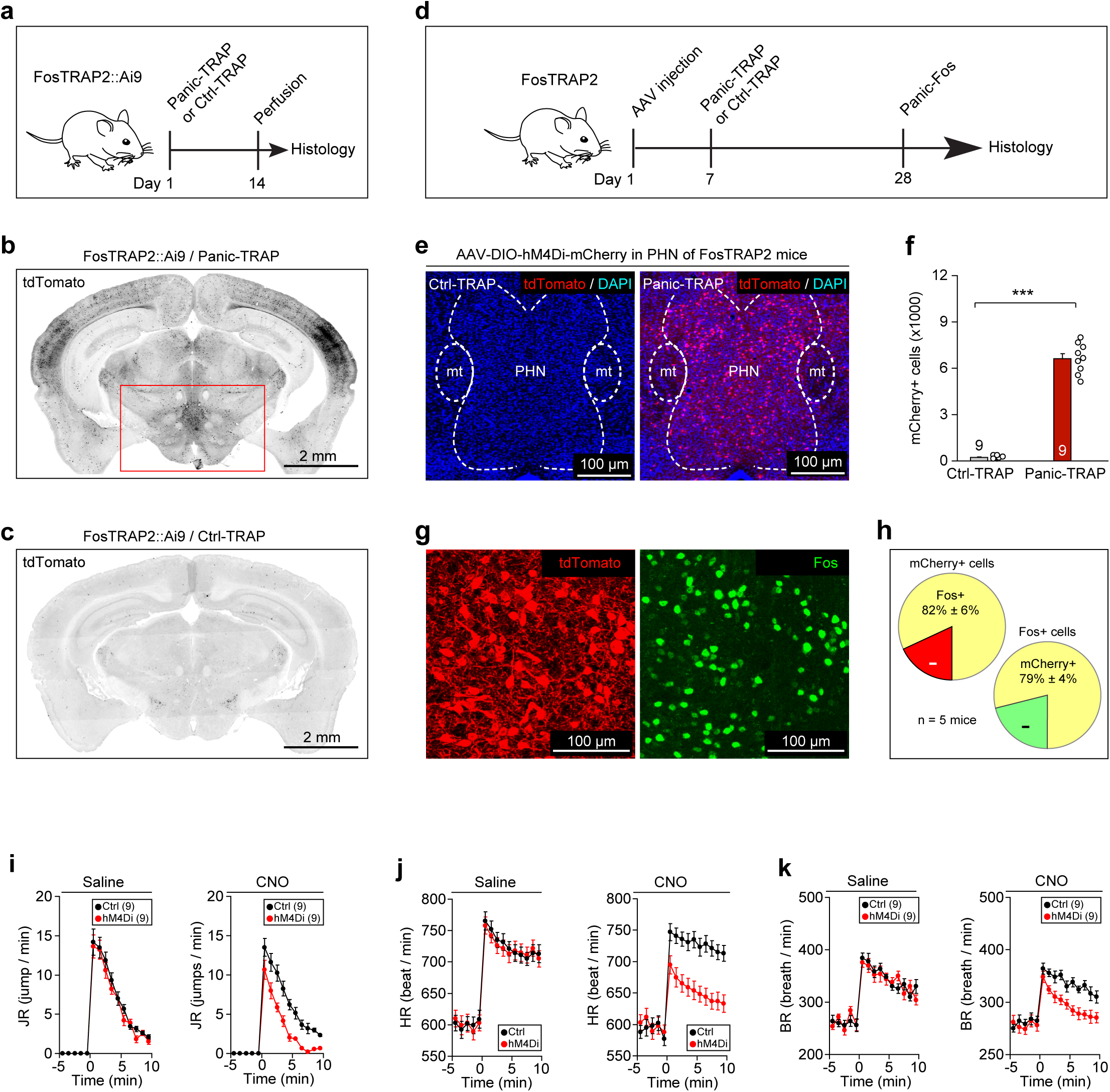
Identification of the PHN as a critical brain area for panic-like defensive state. **(a)** Schematic diagram showing genetic labeling of Panic-TRAP neurons with tdTomato in FosTRAP2::Ai9 mice. Panic-TRAP refers to 4-OH tamoxifen (4-OHT) injection followed by panic induction. Ctrl-TRAP refers to 4-OHT injection without panic induction. **(b, c)** Example coronal brain sections of Panic-TRAP (b) and Ctrl-TRAP (c) mice. **(d)** Schematic diagram showing injection of AAV-DIO-hM4Di-mCherry and subsequent procedure for genetic labeling of Panic-TRAP neurons with hM4Di-mCherry in FosTRAP2 mice. **(e, f)** Example coronal brain sections (e) and quantitative analyses (f) showing efficient labeling of Panic-TRAP neurons in PHN. **(g)** Example coronal brain sections showing colocalization of Panic-TRAP and Panic-Fos cells. **(h)** Quantitative analysis of specificity and efficiency of hM4Di-mCherry for labeling Panic-TRAP PHN neurons. **(i-k)** Time courses of jumping rate (i), heart rate (j), and breathing rate (k) during panic induction in mice treated with saline or CNO (1 mg/kg) to chemogenetically inactivate Panic-TRAP PHN neurons. Scale bars are indicated in graphs. Numbers of mice (f, h, i-k) are indicated in graphs. Data in (f, h, i-k) are means ± SEM. Statistical analyses (f) were performed using Student *t*-test (*** *P* < 0.001). For *P*-values, see Supplementary Table 3.

**Supplementary Fig. 3.**
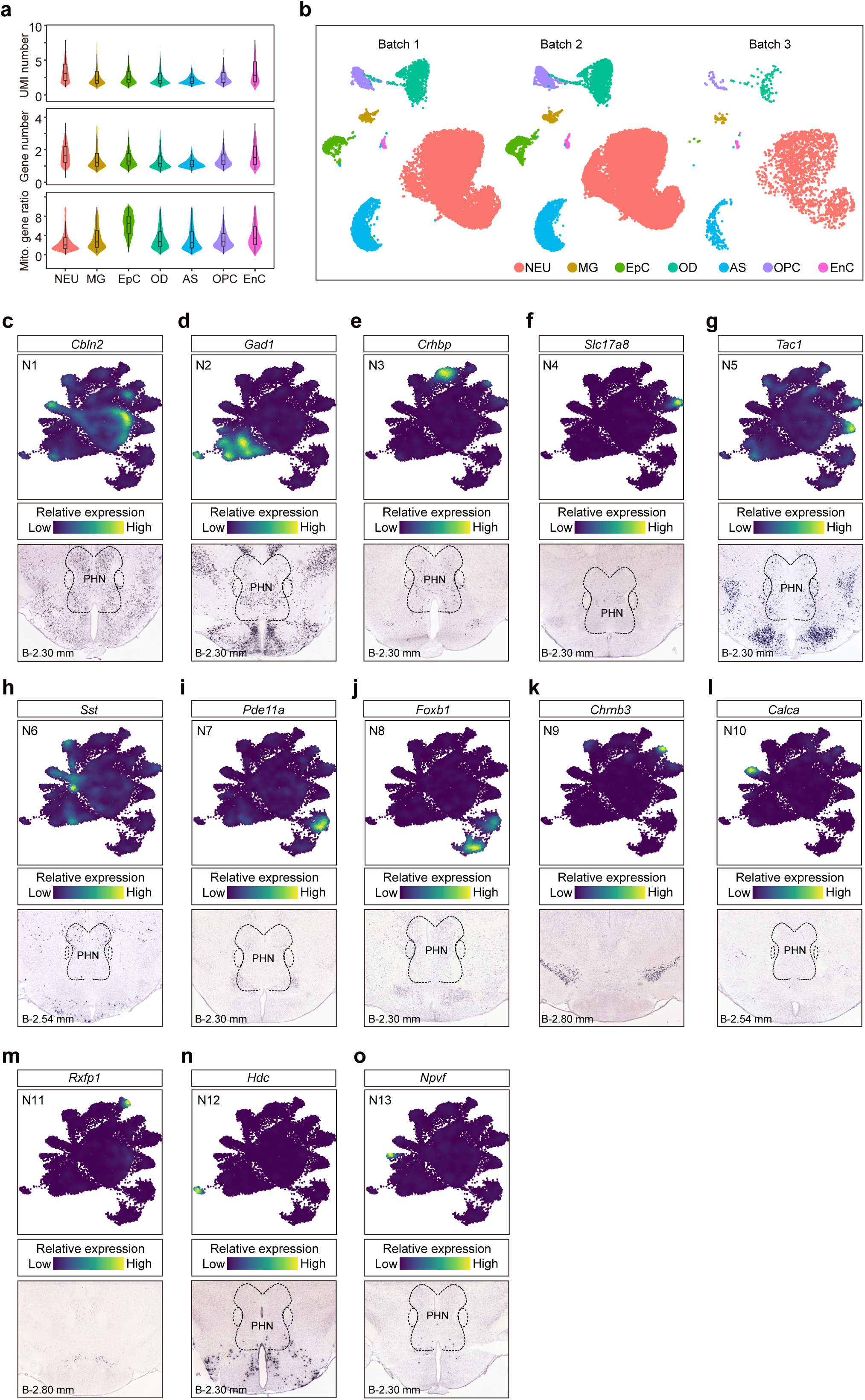
Single-nucleus transcriptomic analysis of cell types in hypothalamic PHN. **(a)** Violin plots showing numbers of unique molecular identifiers (UMIs), detected genes, and mitochondrial gene ratio in each cell class. AS, astrocytes; EnC, endothelial cells; MG, microglia; NEU, neurons; OPC, oligodendrocyte precursor cells; OD, oligodendrocyte cells; EpC, ependymal cells. **(b)** UMAP plot showing sample replicates of snRNA-seq in different batches. **(c-o)** Feature plots showing each neuronal subtype (*top*) and *in situ* hybridization results of marker genes (*bottom*). Note *Cbln2* (c)*, Gad1* (d)*, Crhbp* (e)*, Slc17a8* (f)*, Tac1* (g)*, and Sst* (h) were expressed in PNH, whereas *Pde11a* (i), *Foxb1* (j), *Chrnb3* (k), *Calca* (l), *Rxfp1* (m), *Hdc* (n), and *Npvf* (o) were expressed in brain areas neighboring PHN. Coronal brain sections were adapted from the Allen Mouse Brain Atlas. Abbreviation: *Cbln2, Cerebellin-2; Gad1, Glutamate Decarboxylase 1; Crhbp, Corticotropin Releasing Hormone Binding Protein; Slc17a8, Solute Carrier Family 17 Member 8; Tac1, Tachykinin Precursor 1; Sst, Somatostatin; Pde11a, Phosphodiesterase 11A, Foxb1, Forkhead Box B1; Chrnb3, Cholinergic Receptor Nicotinic Beta 3 Subunit; Calca, Calcitonin Related Polypeptide Alpha; Rxfp1, Relaxin Family Peptide Receptor 1; Hdc, Histidine Decarboxylase; Npvf, Neuropeptide VF Precursor*.

**Supplementary Fig. 4.**
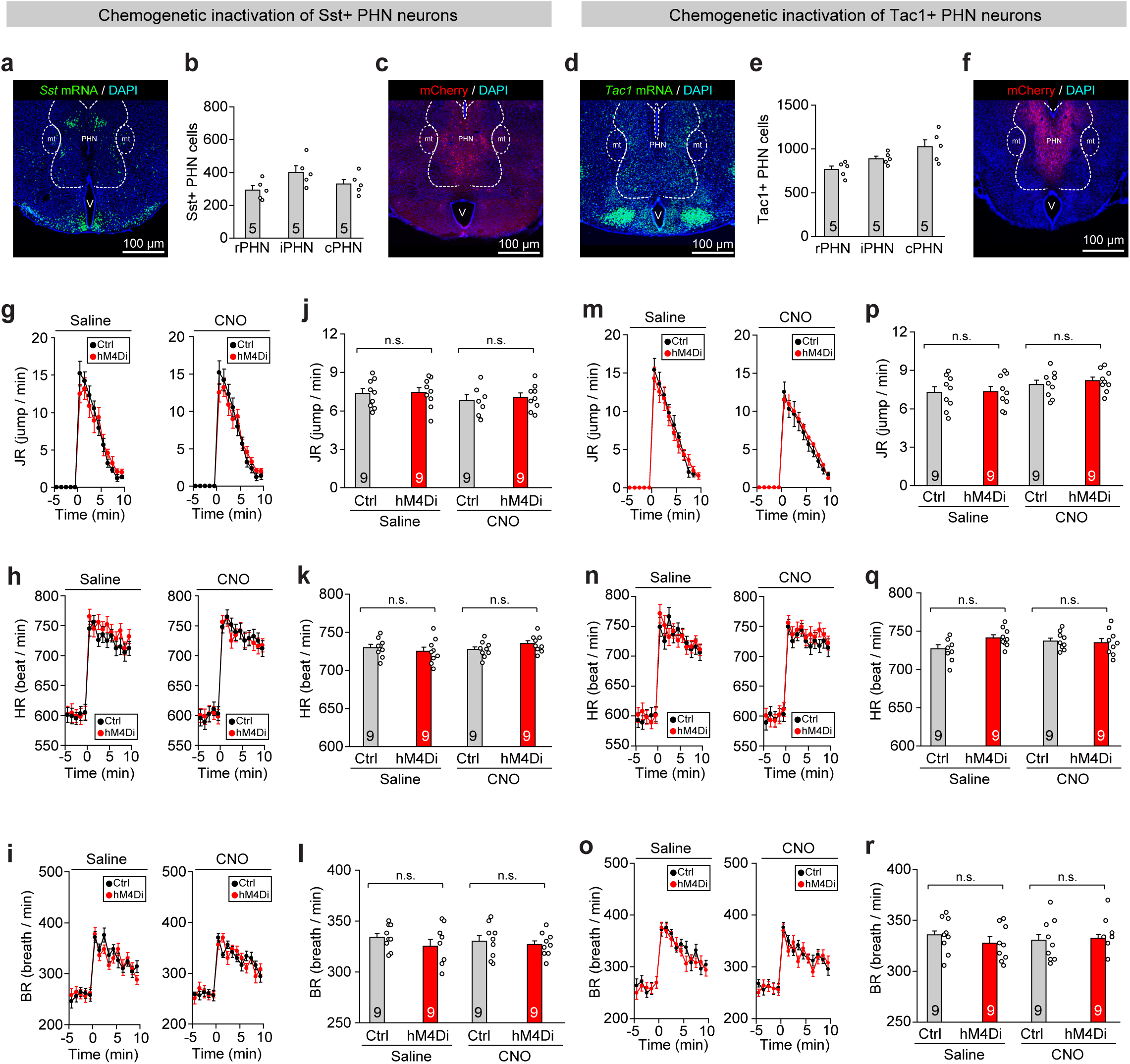
Inactivation of *Sst*+ and *Tac1*+ PHN neurons. **(a, b)** Example coronal section (a) and quantitative analyses (b) showing distribution of *Sst*-expressing (*Sst*+) cells within PHN of WT mouse. **(c)** Example coronal brain section showing hM4Di-mCherry expression in PHN cells of Sst-IRES-Cre mice. **(d, e)** Example coronal section (d) and quantitative analyses (e) showing distribution of *Tac1*-expressing (*Tac1*+) cells within PHN of WT mouse. **(f)** Example coronal brain section showing hM4Di-mCherry expression in PHN neurons of Tac1-IRES-Cre mice. **(g-i)** Time courses of jumping rate (g), heart rate (h), and breathing rate (i) before and during panic induction in mice with and without chemogenetic inactivation of *Sst*+ PHN neurons. **(j-l)** Quantitative analyses of jumping rate (j), heart rate (k), and breathing rate (l) during panic induction in mice with and without chemogenetic inactivation of *Sst*+ PHN neurons. **(m-o)** Time courses of jumping rate (m), heart rate (n), and breathing rate (o) before and during panic induction in mice with and without chemogenetic inactivation of *Tac1*+ PHN neurons. **(p-r)** Quantitative analyses of jumping rate (p), heart rate (q), and breathing rate (r) during panic induction in mice with and without chemogenetic inactivation of *Tac1*+ PHN neurons. Scale bars are indicated in graphs. Numbers of mice (g-r) are indicated in graphs. Data in (g-r) are means ± SEM. Statistical analyses (j-l, p-r) were performed using Student *t*-test (*** *P* < 0.001). For *P*-values, see Supplementary Table 3.

**Supplementary Fig. 5.**
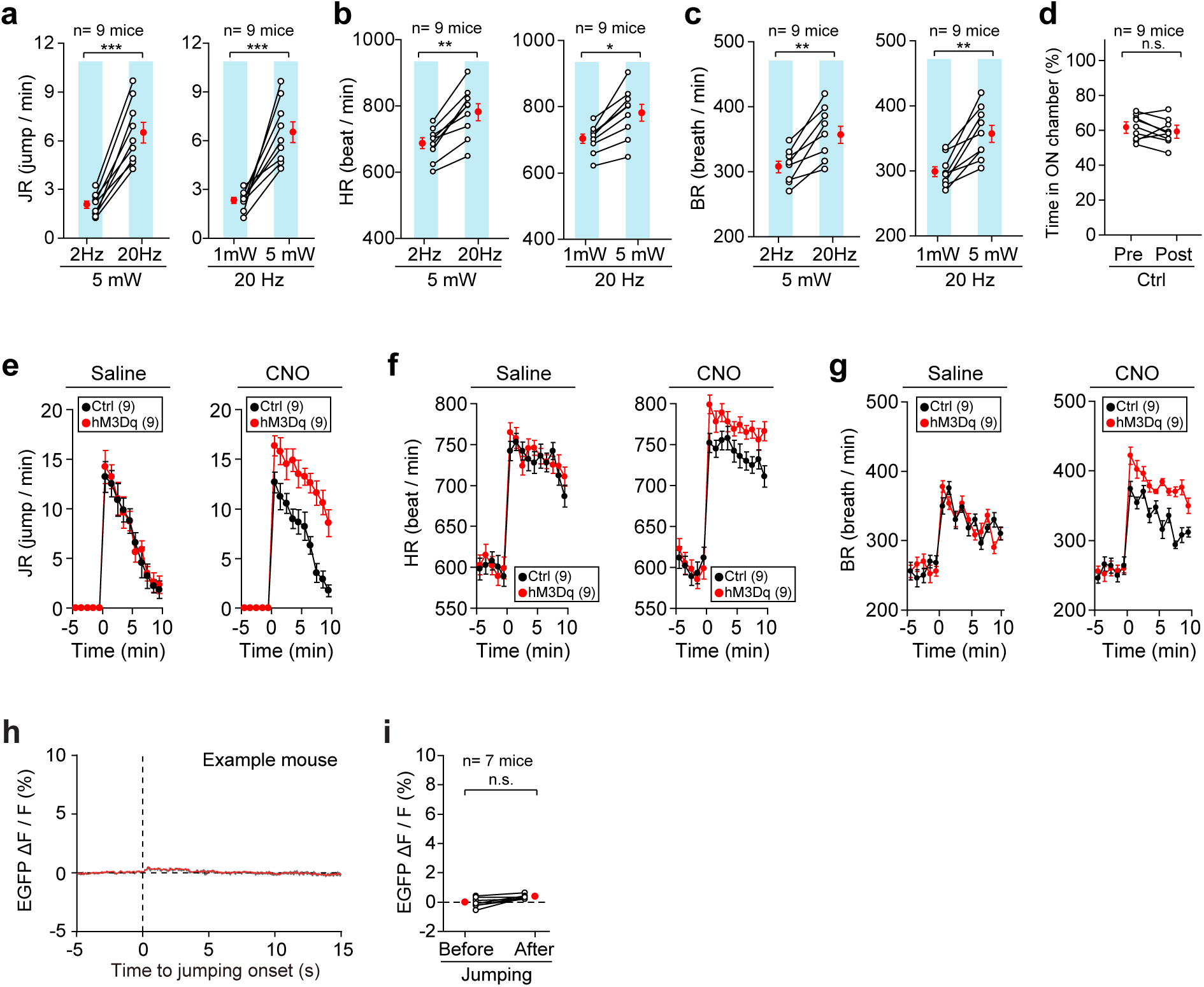
Activation and recording of *Cbln2*+ PHN neurons. **(a-c)** Quantitative analyses of jumping frequency (a), heart rate (b), and breathing rate (c) as a function of light-pulse frequency (left) or laser power (right). **(d)** Quantitative analyses of real-time place aversion in mice with *Cbln2*+ PHN neurons expressing EGFP (Ctrl). **(e-g)** Time courses of jumping frequency (e), heart rate (f), and breathing rate (g) before and during panic induction in mice with and without chemogenetic activation of *Cbln2*+ PHN neurons. **(h)** Average trace of normalized fluorescence changes (ΔF/F) in EGFP-expressing *Cbln2*+ PHN neurons aligned with initiation of jumping of example mouse. **(i)** Quantitative analysis of EGFP fluorescence changes (ΔF/F) in *Cbln2*+ PHN neurons in seven test mice before and after onset of jumping escapes. Numbers of mice (a-g, i) are indicated in graphs. Data in (a-i) are means ± SEM. Statistical analyses (a-d, i) were performed using Student *t*-test (*** *P* < 0.001; ** *P* < 0.01; n.s. *P* > 0.1). For *P*-values, see Supplementary Table 3.

**Supplementary Fig. 6.**
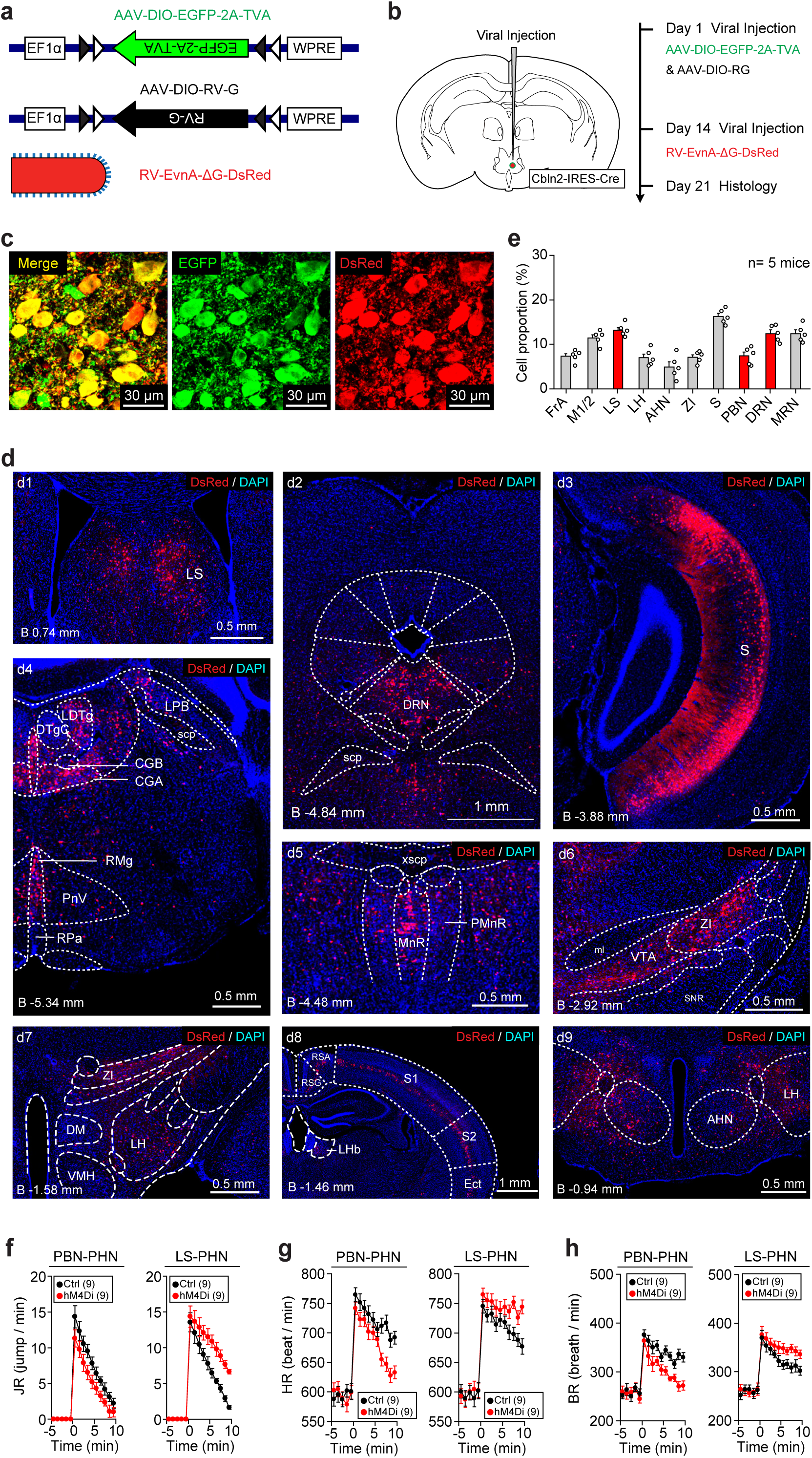
Synaptic inputs of *Cbln2*+ PHN neurons. **(a)** Schematic diagram showing AAV-helpers and RV used in this study. **(b)** Schematic diagram showing timeline for injection of AAV-helpers and RV into PHN of Cbln2-IRES-Cre mice. **(c)** Merged and single-channel example micrographs showing starter cells in PHN. **(d)** Example coronal sections showing distribution of RV-labeled DsRed+ cells in LS (d1), DRN (d2), Subiculum (d3), LDTg (d4), RMg (d4), MnR (d5), ZI (d6), LH (d7), S1/S2 (d8), and AHN (d9). **(e)** Fraction of total RV-labeled DsRed+ cells in selected brain regions monosynaptically projecting to *Cbln2*+ PHN neurons. Data were normalized by dividing total number of DsRed+ cells in these brain regions. **(f-h)** Time courses of jumping rate (f), heart rate (g), and breathing rate (h) before and during panic induction in mice with and without chemogenetic inactivation of PBN-PHN (*left*) and LS-PHN pathways (*right*). Scale bars are indicated in graphs. Numbers of mice (e, f-h) are indicated in graphs. Data in (e, f-h) are means ± SEM.

**Supplementary Fig. 7.**
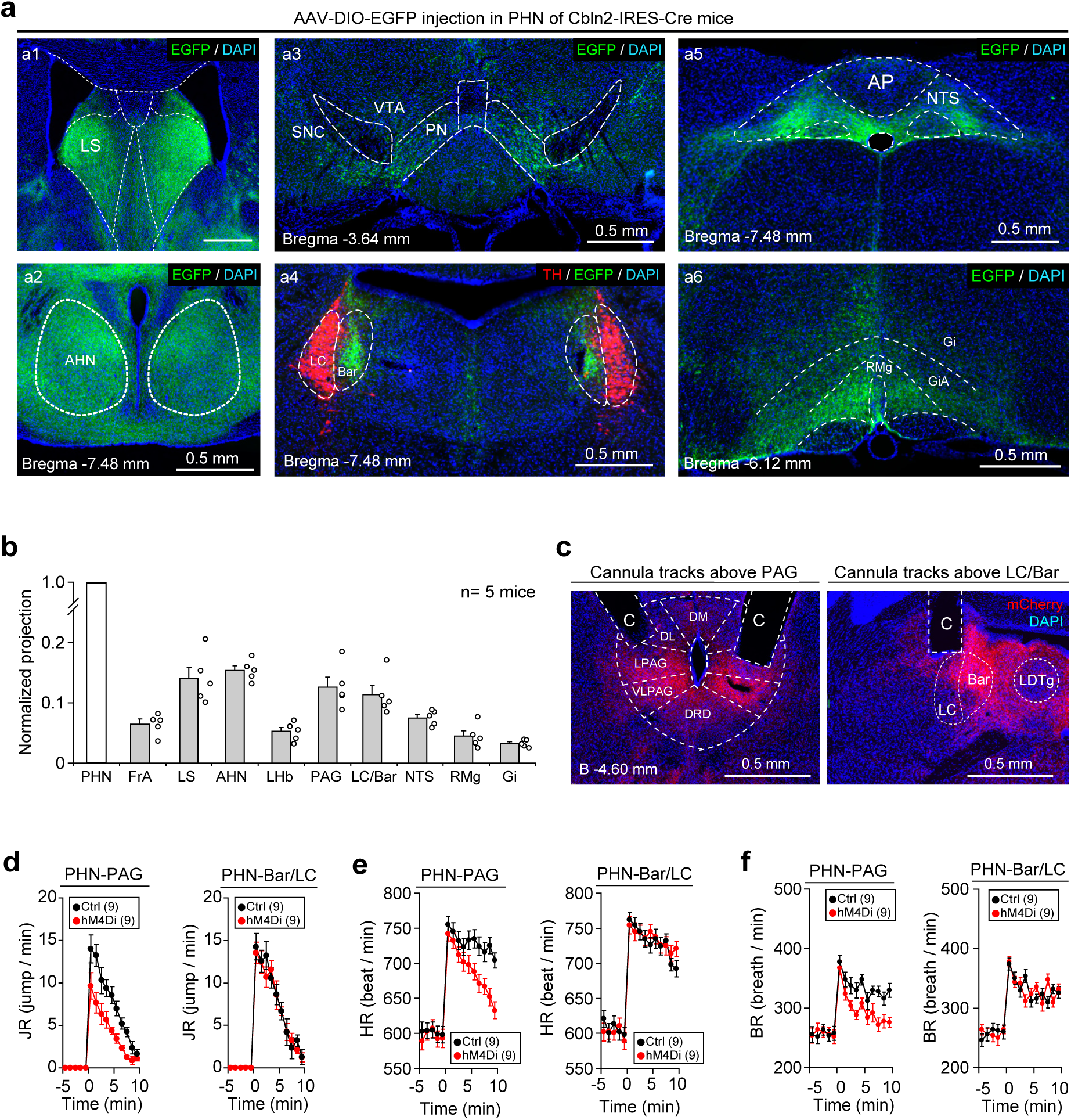
Synaptic outputs of *Cbln2*+ PHN neurons. (a, **b)** Example micrographs (a) and quantitative analyses (b) showing EGFP+ axonal projections of *Cbln2*+ PHN neurons in different brain areas. **(c)** Example coronal sections showing cannula tracks above hM4Di-mCherry+ axon terminals of *Cbln2*+ PHN neurons in PAG (*left*) and Bar/LC (*right*). **(d-f)** Time courses of jumping rate (d), heart rate (e), and breathing rate (f) before and during panic induction in mice with and without chemogenetic inactivation of *Cbln2*+ PHN-PAG (*left*) and PHN-Bar/LC pathways (*right*). Scale bars are indicated in graphs. Numbers of mice (b, d-f) are indicated in graphs. Data in (b, d-f) are means ± SEM.

## SUPPLEMENTARY TABLES

**Supplementary Table 1** Summary of all experimental designs

**Supplementary Table 2** Summary of cell-counting strategies

**Supplementary Table 3** Summary of statistical analyses

**Supplementary Table 4** Differentially expressed genes enriched in each major cluster

**Supplementary Table 5** Differentially expressed genes enriched in each neuronal subtype

**Supplementary Table 6** Reagents, organisms, and instruments

## SUPPLEMENTARY VIDEOS

**Supplementary Video 1**

This video shows robot-induced circa-strike jumping escapes in an example mouse.

**Supplementary Video 2**

This video shows force-plate measurement of jumping escapes in an example mouse.

**Supplementary Video 3**

This video shows robot-induced tachycardia in an example mouse.

**Supplementary Video 4**

This video shows robot-induced tachypnea in an example mouse.

**Supplementary Video 5**

This video shows photostimulation of *Cbln2*+ PHN neurons evoked repetitive jumping escapes in an example mouse.

**Supplementary Video 6**

This video shows GCaMP fluorescence changes in Cbln2+ PHN neurons during panic induction in an example mouse.

**Supplementary Video 7**

This video shows GCaMP fluorescence changes in individual *Cbln2*+ PHN neurons during panic induction of an example mouse.

